# Pharmacological inhibition of PRMT7 links arginine monomethylation to the cellular stress response

**DOI:** 10.1101/503136

**Authors:** Magdalena M Szewczyk, Yoshinori Ishikawa, Shawna Organ, Nozomu Sakai, Fengling Li, Levon Halabelian, Suzanne Ackloo, Mohammad Eram, David Dilworth, Hideto Fukushi, Rachel Harding, Carlo C dela Seña, Tsukasa Sugo, Kozo Hayashi, David Macleod, Carlos Zepeda, Shinji Takagi, Rima Al-Awar, Stephane Richard, Masayuki Takizawa, Cheryl H Arrowsmith, Masoud Vedadi, Peter J Brown, Hiroshi Nara, Dalia Barsyte-Lovejoy

**Affiliations:** Structural Genomics Consortium, University of Toronto, Toronto, Ontario, M5G 1L7, Canada; Research, Takeda Pharmaceutical Company Limited, 26-1, Muraoka-Higashi 2-chome, Fujisawa, Kanagawa 251-8555, Japan; Drug Discovery Program, Ontario Institute for Cancer Research, Toronto, Ontario, Canada; Terry Fox Molecular Oncology Group and Bloomfield Center for Research on Aging, Lady Davis Institute for Medical Research and Departments of Oncology and Medicine, McGill University, Montreal, Quebec H3T 1E2, Canada; Princess Margaret Cancer Centre and Department of Medical Biophysics, University of Toronto, Toronto, Ontario, M5G 2M9, Canada; Department of Pharmacology and Toxicology, University of Toronto, Toronto, Ontario, M5S 1A8, Canada; Nature Research Center, Vilnius, Akademijos 2, Lithuania

**Author notes:** Equal contribution (Yoshinori Ishikawa, Shawna Organ, Nozomu Sakai, Fengling Li). Corresponding authors Dalia Barsyte-Lovejoy, Structural Genomics Consortium, University of Toronto Suite 700, MaRS South Tower, 101 College Street, Toronto, Ontario, M5G 1L7, Canada, Hiroshi Nara, Research, Takeda Pharmaceutical Company Limited, 26-1, Muraoka-Higashi 2-chome, Fujisawa, Kanagawa 251-8555, Japan.

## Abstract

Protein arginine methyltransferases (PRMTs) regulate diverse biological processes and are increasingly being recognized for their potential as drug targets. Here we report the discovery of a potent, selective and cell active chemical probe for PRMT7. SGC3027 is a cell permeable prodrug, which in cells is converted to SGC8158, a potent, SAM-competitive PRMT7 inhibitor. Inhibition or knockout of cellular PRMT7 resulted in drastically reduced levels of arginine monomethylation of HSP70 family members and other stress-associated proteins. Structural and biochemical analysis revealed that PRMT7-driven *in vitro* methylation of HSP70 at R469 requires an ATP-bound, open conformation of HSP70. In cells, SGC3027 inhibited methylation of both constitutive and inducible forms of HSP70, and led to decreased tolerance for perturbations of proteostasis including heat shock and proteasome inhibitors. These results demonstrate a role for PRMT7 and arginine methylation in stress response.

## Introduction

Protein arginine methyl transferases (PRMTs) methylate arginine residues in histone and non-histone proteins in a mono, and symmetric or asymmetric dimethyl manner ^1, 2^. Arginine methylation of both histone and non-histone substrates plays major roles in transcription and chromatin regulation, cell signaling, DNA damage response, RNA and protein metabolism ^3, 4^. PRMT7, a member of the PRMT family, has been functionally implicated in a wide range of cellular processes including DNA damage signaling ^5–7^, imprinting ^8^, and regulation of pluripotency ^9–11^. Recently several elegant studies using *Prmt7* knockout mouse models also revealed the role of this methyltransferase in maintenance of muscle satellite cell quiescence, muscle oxidative metabolism, and B cell biology ^12–14^. While these studies have greatly expanded our understanding of PRMT7 biology, it remains an understudied member of the PRMT family with poor understanding of its cellular substrates.

PRMT enzymes display methylation preference for RGG/RG motifs enriched at protein-protein interfaces, while PRMT7 has been reported to target RXR motifs in arginine and lysine rich regions ^15, 16^. PRMT7 is the sole evolutionary conserved class III PRMT enzyme, the subfamily which carries out only monomethylation of arginine ^17, 18^. Other PRMT family members such as PRMT1 or PRMT5 catalyse arginine dimethylation in an asymmetric or symmetric manner, respectively, playing distinctly different downstream biological roles ^1^. Remarkably, PRMT7-mediated monomethylation of histone H4R17 allosterically potentiates PRMT5 activity on H4R3 19. Thus, possible overlap between substrates for PRMT7 and other PRMT enzymes and their interplay is complex and for most part still largely unknown. The best-characterized PRMT7 substrates are histone proteins, such as H3, H4, H2B and H2A ^1, 3, 6, 18^. Additional non-histone PRMT7 substrates such as DVL3^20^and G3BP2 ^21^ have also been described. Proteomics studies have identified an abundance of cellular monomethyl arginine-containing proteins ^22–25^, however since other PRMT family members may be responsible for this methylation it is not clear which of these substrates are dependent on PRMT7 as systematic studies of PRMT7 cellular substrates are lacking.

To enable further discovery of PRMT7 biology and to better explore its potential as a therapeutic target, we developed a chemical probe of PRMT7 methyltransferase activity. SGC8158 is a potent, selective, and SAM-competitive inhibitor of PRMT7. To achieve cell permeability, we utilized a pro-drug strategy where upon conversion of SGC3027 by cellular reductases, the active component, SGC8158, potently and specifically inhibits PRMT7-driven methylation of cellular substrates. A systematic screen of arginine monomethylated proteins dependent on PRMT7 in cells identified several novel and previously described RG, RGG, and RXR motif proteins. HSP70 family members involved in stress response, apoptosis and proteostasis were further investigated as PRMT7 substrates *in vitro* and in cells. Our data shows that PRMT7 methylates HSPA8 (Hsc70) and HSPA1 (Hsp70) on R469, which resides in a highly conserved sequence in the substrate binding domain. SGC3027 inhibits the PRMT7-driven methylation impacting the thermotolerance and proteostatic stress response in cells.

## Results

### Identification and characterization of selective PRMT7 chemical probe compound

S-adenosyl methionine (SAM) is a co-factor required by all methyltransferases. To identify potential PRMT7 inhibitors, a library of SAM mimetic small molecules was assembled and screened by a scintillation proximity assay (SPA) against PRMT7 using ^3^H-SAM and a histone H2B peptide (residues 23-37) as a substrate (**Supplementary Fig. 1**). This screening resulted in the identification of SGC0911 with IC_50_ of 1 μM (**Fig. 1a**). Further derivatization of this hit yielded SGC8172, which displayed improved potency (IC_50_ <2.5 nM) but was not selective for PRMT7 (**Supplementary Table 1**). Additional optimization of the alkyl chain linking the amine moiety to the adenosine ring resulted in SGC8158 (**Fig. 1a**) which showed good selectivity over a panel of 35 methyltransferases including PRMTs (**Fig. 1b; Supplementary Tables 2 and 3**) while still retaining potent PRMT7 inhibition (IC_50_ <2.5 nM) (**Fig. 1c**). Importantly, we also developed a negative control compound (SGC8158N) which was dramatically less potent (15 ± 2 μM) against PRMT7 (**Fig. 1a,c**) and other protein methyl transferases (**Supplementary Table 2**). Binding of SGC8158 to PRMT7 was also confirmed by surface plasmon resonance (SPR; Supplementary **Fig. 1b,c**). From kinetic fitting, a K_D_ value of 6.4 ± 1.2 nM, k_on_ of 4.4 ± 1.1 × 10^6^ M^-1^ s^-1^ and k_off_ of 2.6 ± 0.5 × 10^-2^ s^-1^ were calculated from triplicate experiments. We next investigated the mechanism of action (MOA) of SGC8158 by determining the IC_50_ values at various concentrations of substrate (SAM and peptide). With increasing concentrations of SAM in the presence of a constant peptide concentration we observed higher IC_50_ values, which indicated a SAM competitive pattern of inhibition (**Fig. 1d**). However, no change in IC_50_ value was observed as the concentration of peptide was increased at fixed concentration of SAM indicating a noncompetitive pattern of inhibition with respect to peptide substrate (**Fig. 1e**).

**Fig. 1:**
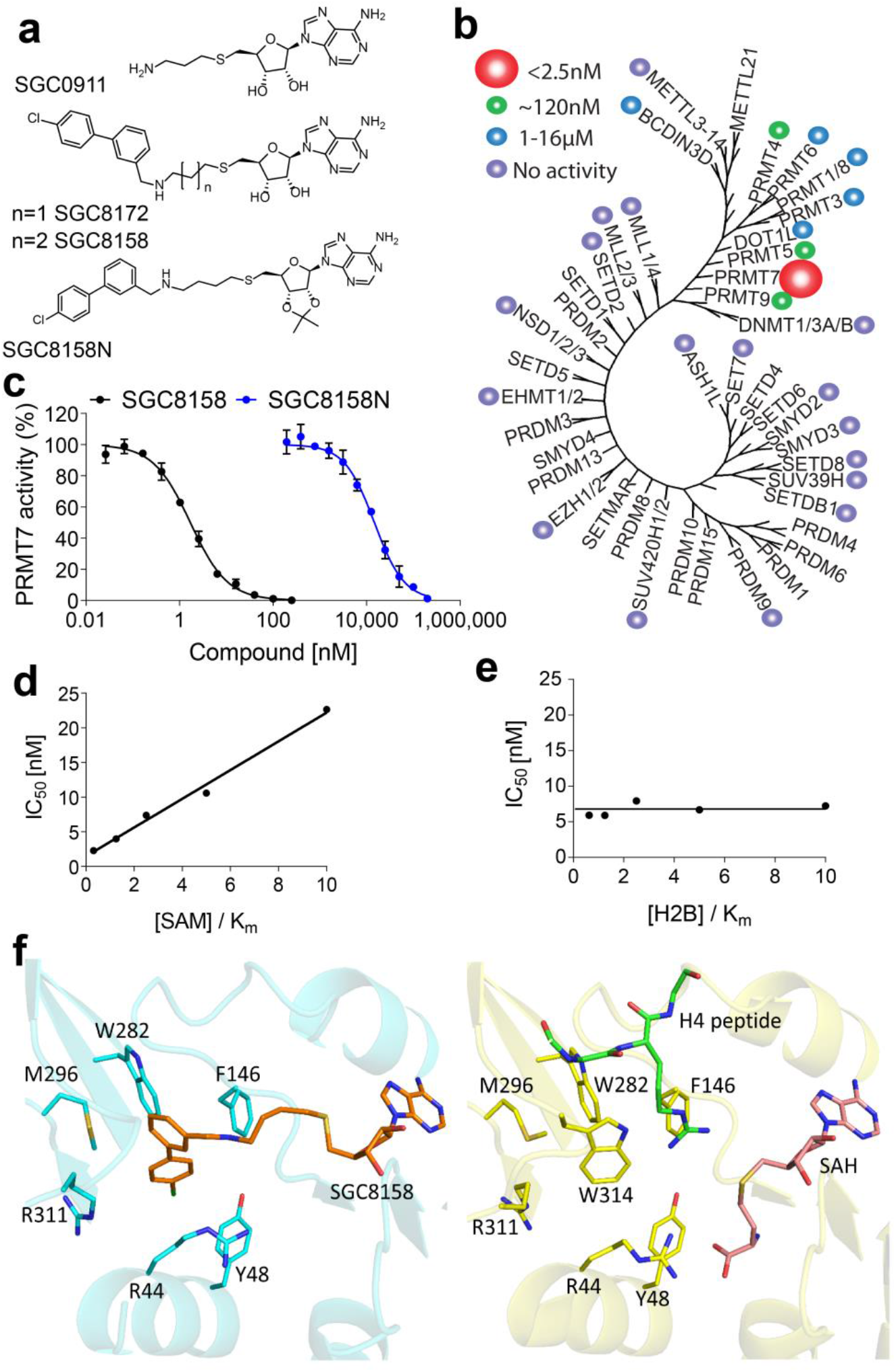
SGC8158 is a potent and selective PRMT7 inhibitor *in vitro*. **a,** Structures of HTS hit compound SGC0911, selective compound SGC8172, active component of the chemical probe SGC8158 and its negative control SGC8158N. **b,** SGC8158 is selective against a panel of 35 protein, DNA and RNA methyltransferases. **c,** SGC8158 inhibits PRMT7 *in vitro* with IC_50_ of < 2.5 nM while negative control compound SGC8158N has IC_50_ of 15 ± 2 μM. The results are Mean+/-SEM, n=3. **d,** SGC8158 is a SAM competitive PRMT7 inhibitor as determined by varying SAM concentrations. **e,** SGC8158 is a peptide non-competitive PRMT7 inhibitor as determined by varying concentration of the peptide H2B. **f,** Crystal structure of mPRMT7 in complex with SGC8158. (Left panel) mPRMT7 is shown as a cartoon representation in cyan, hydrophobic pocket residues are shown in sticks, and the simulated model of SGC8158 is shown in orange. (Right panel) For comparison, the crystal structure of SAH-bound mPRMT7 (PDB ID: 4C4A) is shown as cartoon representation in yellow, hydrophobic pocket residues are shown in sticks, and SAH is shown in pink. Shown in green is the histone H4 peptide from TbPRMT7_H4 (PDB ID: 4M38) superposed onto the mPRMT7-SAH structure.

Human PRMT7 shares 93% sequence identity with *Mus musculus* PRMT7 (mPRMT7), and residues involved in SAM and substrate binding sites are highly conserved between species. To provide further insight into the mechanism of action of SGC8158, we solved the crystal structure of full-length mPRMT7 in complex with SGC8158 refined to 2.94Å resolution, and compared it to that of S-Adenosyl-L-Homocysteine (SAH)-bound mPRMT7 (PDB ID: 4C4A). The ribosyl moiety of SGC8158 is almost identical in position to that of SAH and interacts similarly within the SAM binding pocket, explaining the SAM-competitive kinetics observed in our assays (**Fig. 1f**). Upon soaking mPRMT7-SAH cocrystals with SGC8158, the electron density corresponding to the methionyl moiety of SAH disappeared and new density emerged, leading towards the Arg311-Met315 loop and merging with the electron density of Trp314 side chain (**Supplementary Fig. 2a**). Interestingly, the Arg311-Met315 loop was distorted as a result of soaking with SGC8158, and extra electron density appeared beneath the side chain of Trp314 (**Supplementary Fig. 2a,b**). Accordingly, we reasoned that the biphenylmethylamine moiety of SGC8158 might displace the Trp314 side chain to dock itself into a hydrophobic pocket composed of Trp282, Phe146, Tyr48, Met296, Arg44 and Arg311 residues (**Fig. 1f**). Trp314 is part of the peptide binding pocket that accommodates the Arg3 side chain of histone H4 in peptide-bound *Trypanosoma brucei* PRMT7 (TbPRMT7_H4), (PDB ID: 4M38) (**Fig. 1f**). Comparison of the 311-315 loop region in mPRMT7 with all other PRMTs (**Supplementary Fig. 2c**) showed that this loop has a variable sequence and conformation suggesting that it may play a role in the selectivity of SGC8158 for PRMT7. Taken together, these data indicate that SGC8158 is a potent and selective inhibitor *in vitro* that binds in the adenosine region of the of SAM-binding pocket of PRMT7.

### Identification of novel PRMT7 substrates

In order to evaluate pharmacological inhibition of PRMT7 in cells, we first sought to identify a cellular biomarker of its methylation activity. PRMT7 has a distinct substrate preference for RXR motifs surrounded by basic residues ^17^ and although RGG and RXR motifs are abundant in the proteome ^16^, very few have been validated in the cellular context. The fact that PRMT7 localization is mostly cytoplasmic (**Fig. 2a,b**), and most of the previously investigated substrates are histone proteins (ie. nuclear), led us to undertake a proteomics-based exploration of potential substrates of PRMT7. Wild-type (WT) and *PRMT7* knockout (KO) HCT116 cells were labelled according to the isomethionine methyl-SILAC (iMethyl-SILAC) approach (**Supplementary Fig. 3**)^23^. Of the proteins identified, 280 had at least one monomethylation of arginine and 38 are differentially methylated with a WT/KO fold change equal/greater than 1.3 (**Fig. 2c, Supplementary Table 4**).

**Fig. 2:**
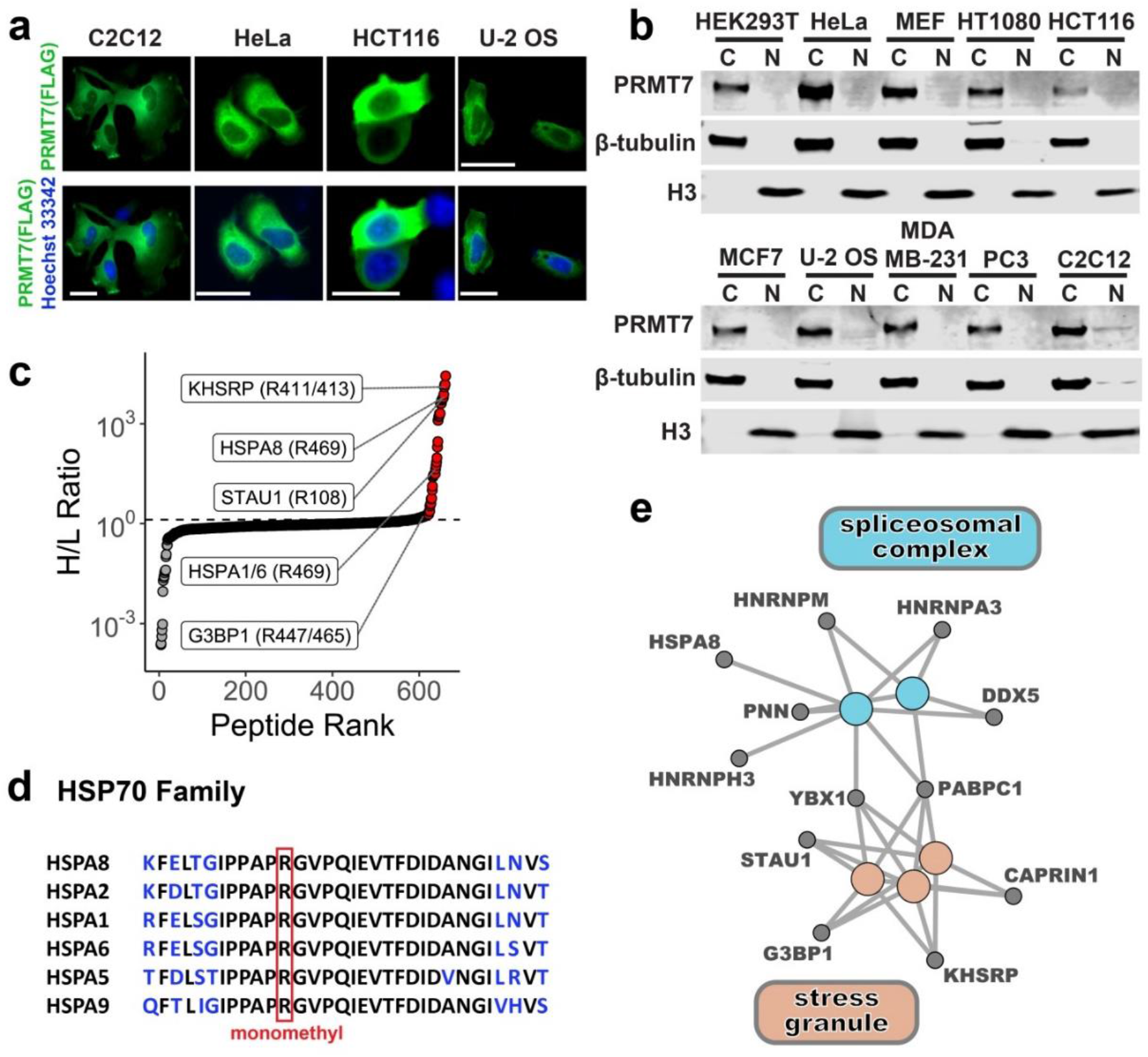
Identification of novel PRMT7 substrates. PRMT7 localizes mainly in the cytoplasm. **a,** Localization of exogenous FLAG-tagged PRMT7 as analysed by IF. Scale bar - 25 μm. Green – FLAG, Blue-Hoechst dye (nucleus). **b,** Cellular fractionation of endogenous PRMT7. C-cytoplasmic fraction, N-nuclear fraction. **c,** Order ranked distribution of heavy (H)/light (L) peptide abundance ratio. The peptides with monomethyl R were identified by MS analysis of HCT116 WT vs *PRMT7* KO cells. Peptides with a H (WT)/L (*PRMT7* KO) ratio equal/greater than 1.3 are indicated in red and the highest ranking peptides for HSPA8, HSPA1/6, STAU and KHSRP are shown. **d,** R469 of HSPA8 resides in a highly conserved region of HSP70 family members. **e,** Network view of enriched cellular component gene ontology terms for proteins with H/L peptide ratios equal/greater than 1.3. Gene ontology terms associated with stress granules are shown in orange and spliceosome complexes in blue, grey nodes represent proteins identified in panel c and are connected to their associated ontologies.

HSPA8 R469 methylation was identified as being highly dependent on PRMT7 and, due to abundant levels of this constitutive HSP70 family member, a commonly used pan monomethyl arginine antibody enabled the detection of this decrease using bulk cell lysates (**Supplementary Fig. 4a,b**). R469 is highly conserved amongst HSP70 family members (**Fig. 2d**) thus in addition to the constitutively expressed HSPA8, HSPA1, or HSPA6, inducible members of the HSP70 family were also identified as putative PRMT7 substrates. Gene pathway analysis of other identified substrates indicated enrichment of proteins associated with RNA metabolism, splicing, and stress granules (**Fig. 2e**).

### PRMT7 methylates HSP70 in cells

In order to confirm that HSPA8 is indeed methylated by PRMT7 in cells, we investigated the endogenous regulation of HSP70 methylation in *PRMT7* KO (HCT116, C2C12 cells), siRNA knockdown (HEK293 and MCF7 cells) and mouse embryonic fibroblasts (MEFs) derived from WT or *Prmt7* KO mice. The *PRMT7* genetic knockout or knockdown resulted in decreased methylation associated with monomethyl arginine signal coinciding with the HSP70 specific signal (**Fig. 3a, Supplementary Fig 4**). As HSP70 proteins are induced and play a role in heat shock response, we investigated if the induced forms of HSP70, such as HSPA6, and HSPA1, are methylated by PRMT7. Heatshock exposure resulted in increased levels of the inducible HSP70 forms, coinciding with increased abundance of the arginine monomethyled species indicating a matched rapid methylation by PRMT7 (**Fig. 3b**).

**Fig. 3:**
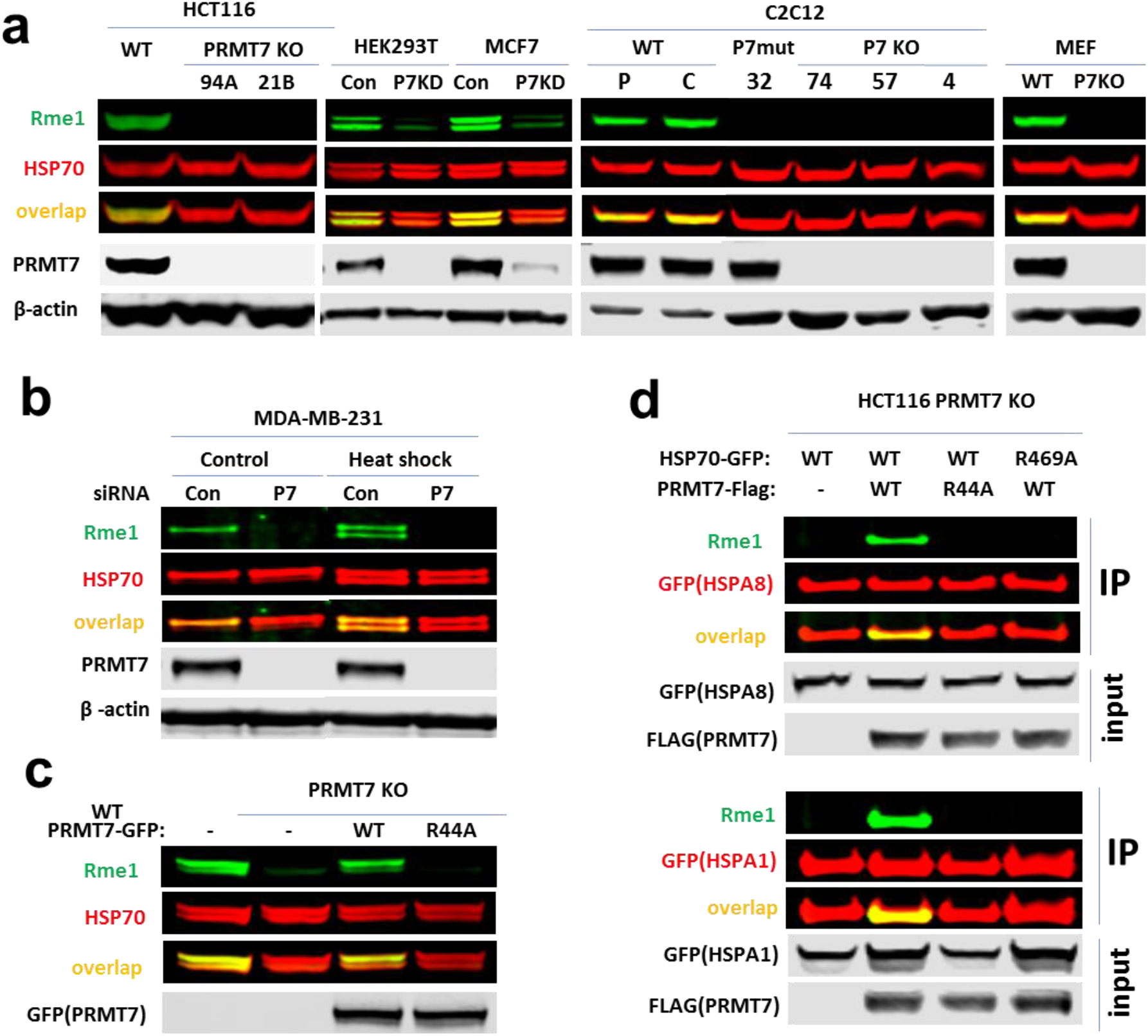
HSP70 R469 is methylated by PRMT7 in cells. **a,** *PRMT7* (P7) knockout (KO) or knockdown (KD) reduces HSP70 methylation in various cell lines. 94A, 21B – HCT116 CRISPR *PRMT7* KO clones; P-parental C2C12; C-C2C12 expressing control guide RNA; 32-C2C12 CRISPR clone expressing PRMT7 catalytic mutant (delY35,A35S); 4,57,74 – C2C12 CRISPR *Prmt7* KO clones. *PRMT7* was knocked down in HEK293T and MCF7 cells using siRNA, Con – control, P7KD – *PRMT7* knockdown. **b,** Monomethylation of inducible and constitutive HSP70 is PRMT7-dependent. MDA-MB-231 cells were transfected with PRMT7 siRNA for 3 days, heat shocked for 1 h at 42 °C and analysed 24 h after heat shock. **c,** Only wild type PRMT7 is able to rescue the HSP70 arginine monomethylation in HCT116 *PRMT7* KO cells. Cells were transfected with GFP-tagged PRMT7 WT or catalytic mutant (R44A). **d,** HSP70 R469A mutation blocks PRMT7 mediated methylation of HSPA8 and HSPA1. HCT116 *PRMT7* KO cells were co-transfected with FLAG tagged *PRMT7* WT or R44A mutant and GFP-tagged HSPA8/1 WT or R469A mutant. HSPA1/8-GFP were immunoprecipitated and analysed for arginine monomethylation levels. The HSP70 methylation in MCF7, HCT116 and HEK293T cells was analysed in cytoplasmic fraction to avoid unspecific band overlap.

The re-expression of WT PRMT7 or the catalytic-dead mutant R44A in HCT116 *PRMT7* KO cells indicated that only WT PRMT7 was able to methylate HSP70 proteins (**Fig. 3c**). Moreover, introduction of WT or R469A mutant HSP70 into HCT116 *PRMT7* KO cells confirmed the methylation site of HSPA8/HSPA1 as identified in the iMethyl-SILAC dataset (**Fig. 3d**). R469 has previously been identified as a methylation site for PRMT4 and PRMT7 ^26^ however, in MCF7 cells, we observed a major effect on HSP70 methylation driven by PRMT7 but not PRMT4 (**Supplementary Fig. 5**) possibly indicating cell-type specific effects. These results demonstrate robust PRMT7-dependent methylation of both steady state and inducible HSP70 proteins in cells.

### PRMT7 methylation of HSP70 *in vitro* reveals a dependency on the open form of HSP70

HSP70 proteins have dynamic structures with nucleotide and substrate binding domains undergoing dramatic conformational changes upon nucleotide or substrate binding ^27, 28^. As R469 resides in the substrate binding domain, we examined the 3D protein structure to determine the accessibility of the R469 side chain. Available full-length structures of HSPA5 (65% sequence identity to HSPA8) and HSPA1A (86% identity) were analysed for the positioning of R469. The structures of HSPA5 in both the ATP-bound open state ^29^ as well as the ADP-bound closed lid state ^29, 30^ for the substrate binding and nucleotide binding domains, respectively, and HSPA1A ^31^ substrate binding domain in the ADP-bound state were used. Analysis of these HSP70 structures indicated that R469 resides in a loop of the substrate binding domain which is likely of limited accessibility in the ADP-bound form of HSP70 when the substrate binding domain is in a closed conformation (**Fig. 4a-c**).

**Fig. 4:**
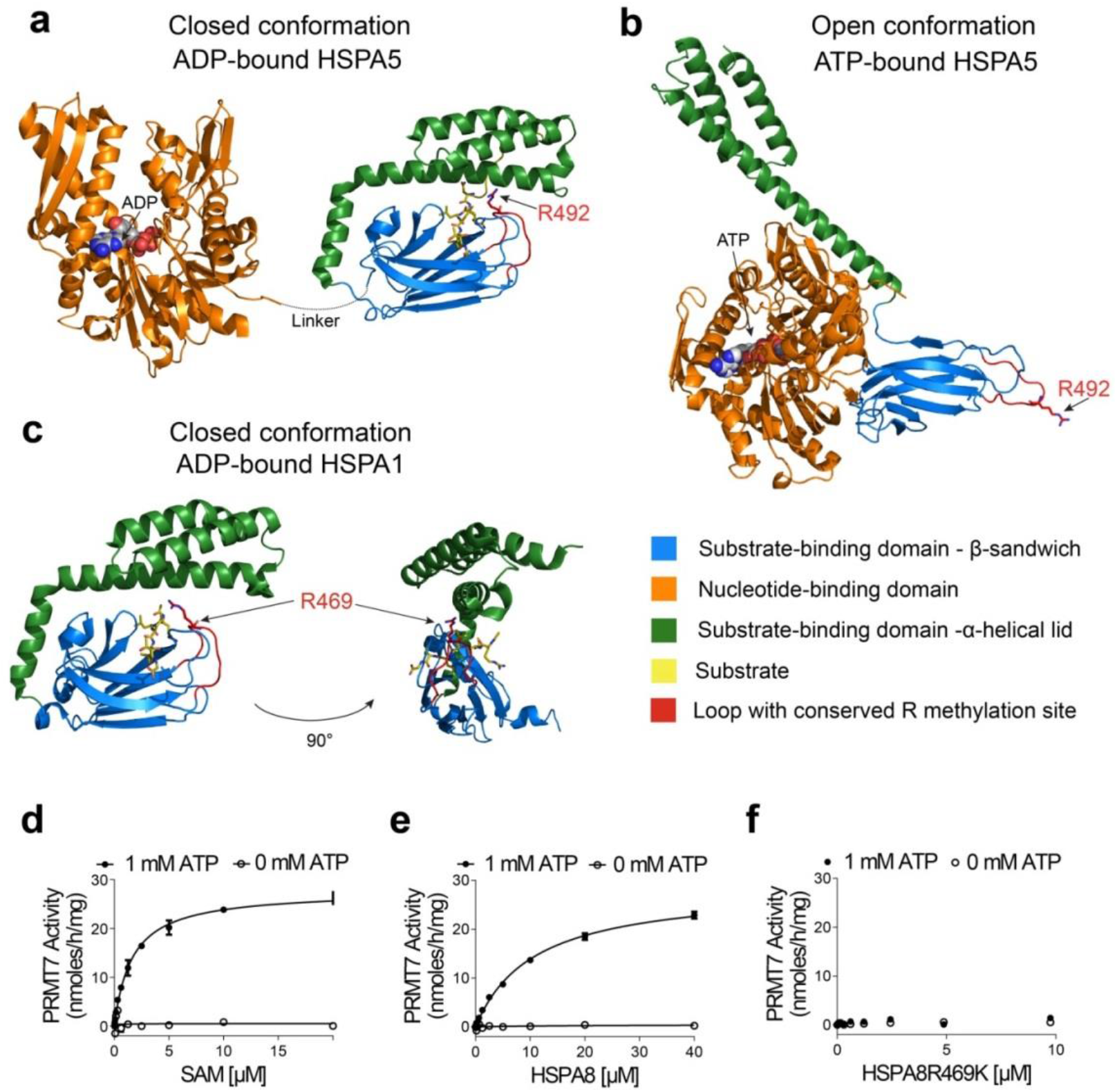
PRMT7 monomethylation of HSP70 depends on the open (ATP-bound) form of HSP70. **a-c,** HSP70 structures in closed and open confirmations reveal differential accessibility of the conserved R469-containing sequence (HSPA8) monomethylated by PRMT7. Structures are color coded for domains and the HSP70 substrate binding domain loop containing PRMT7 methylated arginine is colored red. **a-b,** Closely related homologue HSPA5 structures (65 % overall sequence identity to HSPA8) were analysed to investigate the position of the arginine methylation site in the different conformations. In the ADP-bound state, the lid of the substrate binding domain is closed (PDB 5E85), limiting accessibility of the R492 (analogous to R469 in HSPA8) residue for methylation by PRMT7. In the ATP bound form (PDB 5E84), the arginine residue is accessible therefore permitting access by the PRMT7 enzyme. c, The structure of the more closely HSPA8 related HSPA1A (86 % overall sequence identity and 82 % sequence identity for aa. 386-646 in the substrate binding domain, PDB 4PO2) in the closed conformation in which R469 is occluded by the lid subdomain. **d-f,** Kinetic analysis of HSPA8 methylation by PRMT7 *in vitro*. Kinetic parameters were determined for HSPA8 methylation in the presence and absence of ATP. PRMT7 had no activity in the absence of ATP. **d,** Kinetic analysis at fixed 10 μM HSPA8 (SAM Km = 1.6 ± 0.1 μM). **e,** Kinetic analysis at fixed 20 μM of SAM (HSPA8 Km = 7.2 ± 0.5 μM). **f,** HSPA8 R469K mutant is not methylated by PRMT7 *in vitro*.

In order to establish if PRMT7 methylates HSP70 *in vitro*, we performed full characterization of PRMT7 enzyme kinetics with HSPA8 as substrate. In agreement with structural analysis, our data indicated that PRMT7 methylates HSPA8 in the presence of ATP (**Fig 4d, e**). Apparent kinetic parameters were then determined for HSPA8 methylation yielding a K_m_ for SAM of 1.6 ± 0.1 μM (**Fig 4d**), and K_m_ for HSPA8 of 7.2 ± 0.5 μM (**Fig. 4e**). To test the specificity of this activity, we also tested the activity of PRMT7 on HSPA8 with a single arginine to lysine mutation (R469K). Consistent with previous findings, PRMT7 was completely inactive with the HSPA8-R469K mutant as substrate in the presence or absence of ATP (**Fig. 4f**). We further tested the effect of SGC8158 and SGC8158N on PRMT7-dependent methylation of HSPA8. SGC8158 inhibited PRMT7 methylation of full length HSPA8 *in vitro* with IC_50_ = 294 ± 26 nM under balanced conditions. As expected, SGC8158N showed very poor inhibition with an IC_50_ value estimated to be higher than 100 μM (**Fig. 5a**).

**Fig. 5:**
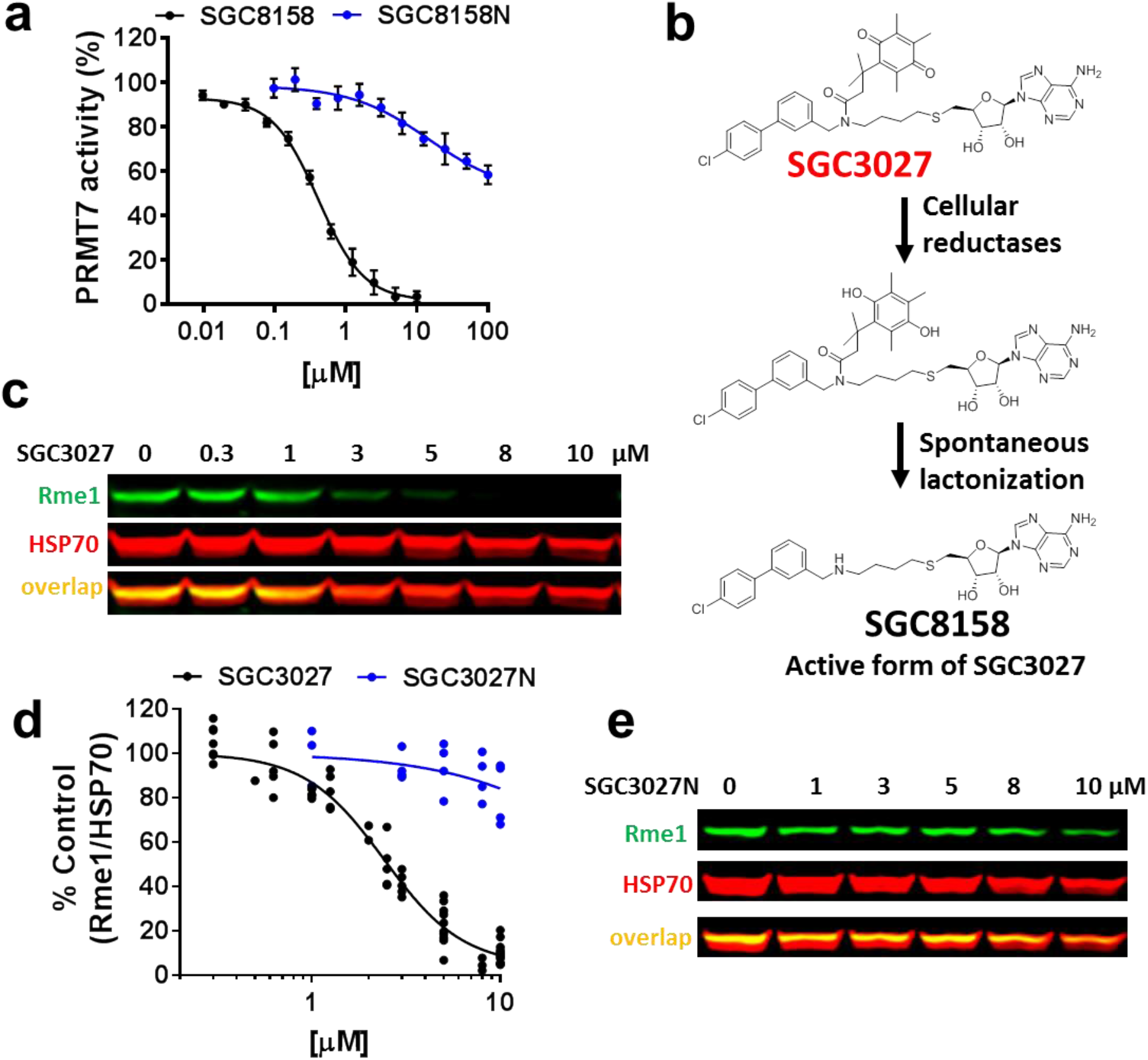
PRMT7 chemical probe SGC3027 inhibits HSP70 methylation in cells and active component SGC8158 inhibits HSPA8 methylation *in vitro*, respectively. **a,** SGC8158 inhibits PRMT7 methylation of HSPA8 *in vitro*. The results are Mean ± SEM (n=3). SGC8158 IC_50_ = 294 ± 26 nM, SGC8158N IC_50_ > 100 μM. **b,** SGC3027 is a prodrug cellular inhibitor of PRMT7 as illustrated by the prodrug conversion to the active component in cells. **c,** SGC3027 inhibits PRMT7 dependent HSP70 monomethylation in C2C12 cells. Cells were treated with the compound for 2 days. **d,** Quantification of SGC3027 and SGC3027N effects on HSP70 monomethylation in C2C12 cells. The graphs represent non-linear fits of Rme1 signal intensities normalized to intensities of HSP70. The results are Mean ± SEM (SGC3027: n=11, 4 separate experiments, IC_50_ = 2.4 ± 0.1 μM; SGC3027N: n=4, IC_50_ > 40 μM). **e,** A representative blot for SGC3027N effects on HSP70 methylation. Rme1 – arginine monomethylation.

### SGC3027, a prodrug form of SGC8158, selectively inhibits PRMT7 in cells

Having established that PRMT7 methylates cellular HSP70, we returned to SGC8158 in order to determine its cellular activity. We observed no inhibition of cellular HSP70 methylation by this compound, likely due to its SAM-like structure and low cell permeability often associated with SAM analogs^32, 33, 34^. To increase the cellular permeability we employed the Trimethyl Lock prodrug strategy in which SGC8158 is derivatized with a quinonebutanoic acid that masks a secondary amine group and increases lipophilicity^35^. The resulting derivative, SGC3027, undergoes reduction in cells followed by rapid lactonization, releasing the active component SGC8158 (**Fig. 5b**).

The SGC3027 prodrug compound inhibited cellular HSP70 methylation with IC_50_ of 2.4 ± 0.1 μM (**Fig. 5c,d**) in C2C12 cells. The inactive compound had no effect at 5 μM, the cellular IC_90_ of SGC3027, and had a minimal effect at 10 μM (**Fig. 5e**). In addition SGC3027, but not SGC3027N, was effective at reducing HSP70 methylation in several commonly used cell lines (**Supplementary Fig. 6**), and SGC3027 was selective over PRMT4, and PRMT5 at 10 μM exposure of (**Supplementary Fig. 7a,b**). In *Prmt7* KO MEFs, SGC3027 does not affect the methylation of HSP70 (**Supplementary Fig. 7c**), further confirming the on-target activity for PRMT7. Taken together these data demonstrate that SGC3027 is a selective and cell-active inhibitor of PRMT7.

### PRMT7 driven methylation is cytoprotective against proteostasis perturbations

HSP70 family members play key roles in nascent protein folding and refolding, as well as distinct roles in anti-apoptotic responses ^36^. To determine if PRMT7-driven methylation contributes to protection from cellular toxic insults, we investigated cell survival in response to thermal and proteasome stress using the genetic *Prmt7* KO models and a chemical biology approach employing the SGC3027 chemical probe. *Prmt7* KO MEF cells were more sensitive to acute heat stress with fewer cells surviving the treatment (**Fig. 6a**) and more cells undergoing apoptosis (**Fig. 6b**). SGC3027, but not the negative control SGC3027N, showed a similar response (**Fig. 6c,d**), indicating that SGC3027 phenocopies the genetic ablation of *Prmt7*. We also investigated if PRMT7 stress protection extends to proteasome stress using proteasome inhibitor bortezomib. Compared to WT cells, *Prmt7* KO MEFs cells were more sensitive to acute bortezomib induced cell death, and had poorer recovery after 4 or 20 h exposure to bortezomib (**Fig. 6e,f**). SGC3027, but not SGC3027N, sensitized the WT *Prmt7* MEFs to bortezomib while neither compound affected *Prmt7* KO cells (**Fig. 6g,h**). These results indicate that PRMT7-driven methylation plays a cytoprotective role in stress response, while inhibition of PRMT7 catalytic function can sensitize cells to toxic stimuli.

**Fig. 6:**
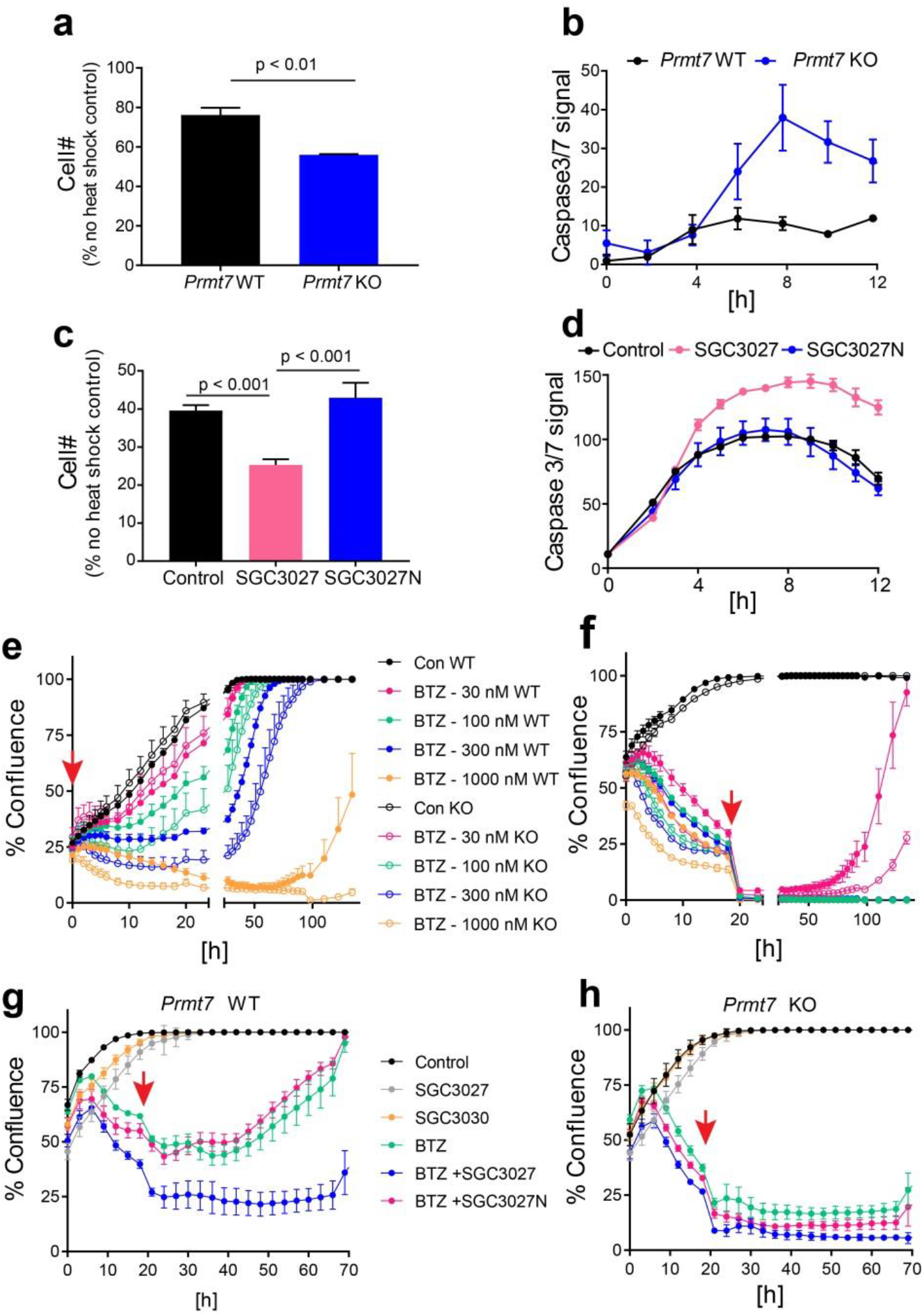
PRMT7 knockout/inhibition affects cell survival after heat shock or proteasomal stress. **a-b,** Loss of PRMT7 decreases cell survival and increases apoptosis levels after heat shock. MEF cells were heat shocked for 20 min at 44 °C. Apoptosis was monitored immediately after the heat shock and cell number was determined 24 h later. The results are Mean ± SEM of 3 replicates. **c-d,** SGC3027 inhibition of PRMT7 activity decreases cell survival and increases apoptosis levels after heat shock. Cells were pretreated with 3 μM compounds for 2 days before heat shock. Apoptosis and cell number were determined as in **a** and **b**. The results are Mean ± SEM of 3 replicates. **e-f,** Loss of PRMT7 decreases cell survival after bortezomib (BTZ) treatment. BTZ was removed after 4 h (**e**) or 24 h (**f**) and the cell confluence was monitored 4 h after BTZ treatment. The results are Mean ± SEM of 6 replicates. **h-i,** SGC3027 decreases cell survival after BTZ treatment (30 nM) in *Prmt7* WT MEF (**g**) but not in *Prmt7* KO MEF (**h**). Cells were pretreated with 4 μM compounds for 2 days before BTZ treatment. After 20 h, BTZ was removed and SGC3027 or SGC3027N were replaced. The results are Mean ± SD of 4 replicates. Red arrow indicates the time BTZ was removed.

## Discussion

PRMT7 is a monomethyl arginine methyltransferase that plays a role in muscle physiology and stem cell biology ^3, 10–13^. However, the PRMT7 substrates that mediate this biology are not well understood, and tools to selectively and temporally modulate PRMT7 catalytic activity are lacking. Here we report the first selective PRMT7 chemical probe, SGC3027, and a closely related but inactive compound, SGC3027N, for use as a specificity control for biological experiments. We also identified novel PRMT7 methylation substrates using proteomics and further validated HSPA8 and related HSP70 family members as PRMT7 substrates by employing *in vitro* assays, genetic methods, and chemical biology. Our data suggest that PRMT7 activity plays a role in cellular thermotolerance and proteasomal stress response. Therefore, SGC3027 maybe be a useful modulator of cellular proteostasis in stress response under physiological and pathological conditions.

SGC8158 is a structural derivative of SAM, acts as a SAM-competitive inhibitor of PRMT7, and occupies the adenosine pocket in the SAM binding site of PRMT7. Other potent and selective SAM-like inhibitors of methyltransferases including EPZ004777 ^37^ and SGC0946 for DOT1L ^38^ and LLY-283 for PRMT5 ^39^ feature extensively modified adenosine and methionine moieties that likely enhance the cell permeability of otherwise cell impermeable SAM. We employed an alternative prodrug strategy in which adding the quinonebutanoic moiety increased cell permeability and allowed for cellular reductases to generate the active component, SGC8158. Notably, the cellular reductase-driven activation of SGC3027 to the active SGC8158 may vary among cell types depending on the abundance or activity of reductase enzymes. Thus, we recommend that the appropriate concentration range for use of these probes should be empirically evaluated in each experimental setting by monitoring a biomarker such as HSP70 methylation before evaluation of specific biological readouts.

SGC3027 sensitized cells to heat and proteasomal stresses indicating that PRMT7 catalytic function is required for normal physiological response to these stimuli. *Prmt7* KO cells were also more sensitive to these stresses, indicating that the chemical probe phenocopies the genetic knockout effects. Heat stress and proteasome inhibition elicits orchestrated protective responses, with the central goal of maintenance of cell protein homeostasis, proteostasis. Failure to maintain proteostasis results in numerous diseases. For example, altered proteostatic balance due to rewiring of chaperome complexes has been observed in cancer cells, contributing to drug resistance ^40, 41^. PRMT7-dependent protection against cellular stress may have physiological importance in cancer cell survival, consistent with higher levels of PRMT7 that have been reported in breast cancer cell lines ^42, 43^. Interestingly, *PRMT7* was initially identified in a screen for sensitization to topoisomerase inhibitors in cancer cells ^5^, suggesting a wider range of stressors against which PRMT7 may be protective. Our findings of PRMT7 inhibition leading to the sensitization of cells to bortezomib-induced cell death indicate potential therapeutic applications in cancers such as multiple myeloma and some lymphomas. It is possible that PRMT7 inhibition may lead to overcoming resistance to bortezomib or other therapies. Further work is needed to determine if PRMT7 inhibition can indeed synergize with proteasome inhibitor drugs in therapeutically relevant cell systems, and these efforts will be aided by the use of SGC3027 as a chemical probe for PRMT7.

The majority of PRMT7 substrates identified in this study were functionally associated with stress response, RNA metabolism, splicing, and stress granules. Notably G3BP2, a previously reported PRMT7 substrate ^21^, STAU1 ^44^ and KHSRP ^45^ have all been linked to stress granule biology and cell stress response. Previous arginine methylome studies have also noted the abundance of mono and dimethyl arginine posttranslational modifications (PTMs) in proteins involved in splicing, transcription, and RNA metabolism ^22–25, 46, 47^. We identified and validated members of the HSP70 family as cellular substrates of PRMT7 leading us to further investigate the role of PRMT7 in stress response.

The cytoprotective function of HSP70 proteins has been attributed to the modulation of protein refolding and transport of client proteins, as well as direct regulation of apoptotic signaling pathways ^40, 48^. Several HSP70 inhibitors have been reported and their utility is being explored in counteracting bortezomib resistance in multiple myeloma as well as therapeutic applications in other cancers ^49–52^. PTMs of the HSP70 family members such as lysine methylation, ubiquitination, acetylation, and phosphorylation have been reported ^53^ and some of the PTMs stabilize the HSP70/HSP90/HSP40 and client protein complexes allowing the formation of antiparallel HSP70 dimers ^54^. K561 trimethylation by METTL21A and dimethylation by SETD1A have been reported to result in the modulation of HSP70 affinity for client proteins or potentiation of AURKB activity, respectively ^55–57^. Here, using genetic and pharmacological means, we identify R469 monomethylation as an abundant cellular modification that depends on PRMT7 catalytic function and correlates with the cytoprotective properties of PRMT7.

HSP70 R469 monomethylation by PRMT4 and PRMT7 was recently reported as contributing to transcriptional activation by retinoic acid receptor RAR ^26^. Interestingly PRMT4 methylation of HSP70 did not seem to require an ATP-driven conformational change ^26^ while we found that PRMT7 driven methylation occurs when HSP70 adopts an open, ATP-bound conformation that promotes R469 accessibility. In all of the examined structures of the substrate and ADP-bound forms of HSP70, R469 packs into the lid subdomain of the substrate binding domain to ensure flexible, yet stable, interaction with the client protein ^31^, but rendering this residue inaccessible to modifying enzymes such as PRMT7. Although the substrate binding domains and lid subdomains are more variable among HSP70 proteins than other regions, possibly due to the need of accommodating a large number of substrates ^31^, the region surrounding R469 is highly conserved among HSP70 family members and species. PTMs of the HSP70 lid and tail domains have been reported to disrupt the interaction with co-chaperone CHIP TPR ^58^. Since the R469-containing loop resides in close proximity to the CHIP binding region it is also possible that the methylation may influence HSP70 co-chaperone recruitment. Further experiments, required to completely understand the mechanistic role of R469 methylation in HSP70 function, will benefit from the use of this PRMT7 chemical probe.

We uncovered new aspects of PRMT7 biology associated with proteostasis, identified PRMT7 cellular substrates and validated HSP70 family members HSPA1/6/8 as PRMT7 substrates, whose methylation is likely to contribute to the cytoprotective effects of PRMT7. SGC3027, together with its negative control SGC3027N, will be useful tools for further understanding PRMT7 function in physiological and disease states.

## Acknowledgements

The SGC is a registered charity (number 1097737) that receives funds from AbbVie, Bayer Pharma AG, Boehringer Ingelheim, Canada Foundation for Innovation, Eshelman Institute for Innovation, Genome Canada through Ontario Genomics Institute [OGI-055], Innovative Medicines Initiative (EU/EFPIA) [ULTRA-DD grant no. 115766], Janssen, Merck KGaA, Darmstadt, Germany, MSD, Novartis Pharma AG, Ontario Ministry of Research, Innovation and Science (MRIS), Pfizer, São Paulo Research Foundation-FAPESP, Takeda, and Wellcome [106169/ZZ14/Z].

This work is based upon research conducted at the Northeastern Collaborative Access Team beamlines, which are funded by the National Institute of General Medical Sciences from the National Institutes of Health (P30 GM124165). The Eiger 16M detector on 24-ID-E beam line is funded by a NIH-ORIP HEI grant (S10OD021527). This research used resources of the Advanced Photon Source, a U.S. Department of Energy (DOE) Office of Science User Facility operated for the DOE Office of Science by Argonne National Laboratory under Contract No. DE-AC02-06CH11357.

We thank Kumar Singh Saikatendu, Charles E. Grimshaw, Nobuo Cho, Toshiyuki Nomura, and Atsushi Kiba for suggestions and support throughout the project and Michiko Tawada and Sachiko Itono for suggestions and chemistry support. We thank Alma Seitova, Ashley Hutchinson, Hong Zheng for recombinant protein production and Aiping Dong for crystallography support.

## Methods

### Protein expression and purification

Full-length PRMT7 was expressed in Sf9 cells grown in HyQ® SFX Insect serum-free medium (ThermoScientific). Cells were harvested and lysed, and the cleared lysate was incubated with 5 mL anti-FLAG M2-Agarose (Sigma) in 50 mM Tris-HCl, pH 7.5, 150 mM NaCl, 10% glycerol and washed with the same buffer with 500 mM NaCl. The pure recombinant protein was eluted from the column using the same buffer with 0.1 mg.mL^-1^ FLAG peptide (Sigma). Pure PRMT7 was flash frozen and stored at −80 °C. Full length HSPA8 was overexpressed in *Escherichia coli* strain BL21(DE3) V2R-pRARE2 during an overnight induction with 0.5 mM isopropyl 1-thio-d-galactopyranoside at 18 °C. Cells were suspended in 20 mM Tris-HCl (pH 8.0, 1 mM DTT, 300 mM NaCl). The clarified lysate was loaded onto a Hispur™ nickel-nitrilotriacetic acid column (Thermo Scientific) and washed with buffer. Then, protein was eluted and concentrated. Protein purity was determined by SDS-PAGE and LC-MS.

### Radioactive activity assay *in vitro*

Assays using biotinylated H2B (23-37) as a substrate were performed in buffer (20 mM Tris-HCl, pH 8.5, 0.01% Tween-20, and 5 mM DTT) containing 5 nM PRMT7, 1.1 μM ^3^H-SAM (Cat.# NET155V250UC; Perkin Elmer; www.perkinelmer.com) and 0.3 μM H2B (23-37). The reaction mixtures were incubated for 60 min at 23 °C. To stop the enzymatic reactions, 10 μL of 7.5 M guanidine hydrochloride was added, followed by 180 μL of buffer (20 mM Tris, pH 8.0), mixed and then transferred to a 96-well FlashPlate (Cat.# SMP103; Perkin Elmer; www.perkinelmer.com). After mixing, the reaction mixtures in Flash plates were incubated for 1 hour and the CPM were measured using Topcount plate reader (Perkin Elmer, www.perkinelmer.com). The CPM counts in the absence of compound for each data set were defined as 100% activity. In the absence of the enzyme, the CPM counts in each data set were defined as background (0%). The IC_50_ values were determined using GraphPad Prism 7 software. For the kinetic analysis of HSPA8 methylation by PRMT7, the assay mixture contained 20 mM Tris-HCl, pH 8.5, 0.01% Tween-20, and 5 mM dithiothreitol (DTT), 1 mM MgCl_2_, 1 mM ATP where indicated, 250 nM PRMT7, fixed concentration (20 μM) of SAM, various concentrations (up to 40 μM) of HSPA8; or fixed concentration (10 μM) of HSPA8 with different concentrations of SAM (up to 20 μM). IC_50_ determinations of SGC8158 and SGC8158N were performed at 150 nM PRMT7, close to K_m_ values of both SAM (2 μM) and substrate HSPA8 (7 μM). To determine the mode of action, the experiments were performed in the presence of fixed biotinylated H2B peptide (residues 23-37) or HSPA8 substrate concentration and increasing SAM concentration or at fixed concentration of SAM and varying concentration of the substrate. Twenty μL of reaction mixtures were incubated at 23 °C for 60 min. To stop reactions, 100 μL of 10% trichloroacetic acid (TCA) was added, mixed and transferred to filter-plates (Millipore; cat.# MSFBN6B10; www.millipore.com). Plates were centrifuged at 2000 rpm (Allegra X-15R - Beckman Coulter, Inc.) for 2 min followed by 2 additional 10% TCA washes and one ethanol wash followed by centrifugation. Plates were dried and 30 μL MicroScint-O (Perkin Elmer) was added to each well, centrifuged and removed. 50 μL of MicroScint-O was added again and CPM was measured using Topcount plate reader. The IC_50_ values were determined using GraphPad Prism 7 software.

### SPR analysis

SPR analysis was performed using a Biacore™ T200 (GE Health Sciences Inc.) at 20 °C. Approximately 5500 response units of Bio-PRMT7 (amino acids 1-692) was fixed on a flow cell of a SA chip according to manufacturer’s protocol, while another flow cell was left empty for reference subtraction. SPR analysis was performed in HBS-EP (20 mM HEPES pH 7.4, 150 mM NaCl, 3 mM EDTA, 0.05% Tween-20) with 3% DMSO. Five concentrations of SGC8158 (150, 50, 16.6, 5.5 and 1.85 nM) were prepared by serial dilution. Kinetic analysis was performed using single cycle kinetics with contact time of 60 sec, off time of 300 sec, and a flow rate of 100 μL.min^-1^. To favor complete dissociation of compound for the next cycle, a regeneration step (300 second, 40 μL.min^-1^ of buffer), a stabilization period (120 sec) and two blank cycles were run between each cycle. Kinetic curves were fitted using a 1:1 binding model and the Biacore T200 Evaluation software (GE Health Sciences Inc.).

### Selectivity assays

The methyltransferase selectivity was assessed at two compound concentrations of 1 and 10 μM as described previously ^59–61^.

### Constructs, Cells and Antibodies

MCF7 (ATCC® HTB-22™), C2C12, MEF WT and MEF *Prmt7KO* (kind gift from Dr. Stephane Richard, McGill University), U-2 Os (ATCC®HTB-96™), HT-1080 (ATCC®CCL-121™) and HEK293T (kind gift from Sam Benchimol, York University) were grown in DMEM (Wisent), HCT116 WT (ATCC®CCL-247™), in McCoy’s (Gibco) and MDA-MB-231 (ATCC® HTB-26™) in RPMI1640 (Wisent) supplemented with 10% FBS (Wisent), penicillin (100 U.mL^-1^) and streptomycin (100 μg.mL^-1^). Anti-Rme1 (#8015), anti-Rme2s (#13222) and anti-mouse Alexa Fluor 488 (#4408) were purchased from Cell Signaling Technologies. Anti-Hsp/Hsc70 was from Enzo (#ADI-SPA-820). Antibodies for PRTMT7 (#ab179822), PRMT5 (#ab109451), and β-actin (#ab3280) were purchased form Abcam. Anti-PRMT4 (#A300-421A) was from Bethyl. Anti-GFP (#632381) used for western blot was purchased from Clontech. Anti-GFP used for IP was purchased from Invitrogen (#G10362). Anti-Flag (#F4799) was from Sigma. Anti-SmBB’ (#sc-130670) and anti-BAF155 (#sc-32763) was from Santa Cruz Biotechnologies. Anti-BAF155-R1064me2a (#ABE1339) was from Millipore. Goat-anti rabbit-IR800 (#926-32211) and donkey anti-mouse-IR680 (#926-68072) were purchased from LiCor. Full length of HSPA8, HSPA1 were cloned into pAcGFPN3 vector (Clontech) and PRMT7 were cloned into pAcGFPN3 (Clontech) or pcDNA3 (N-terminus FLAG). Site directed mutagenesis to generate PRMT7 R44A mutant, HSPA1 R469A and HSP8 R469A mutants was performed using Q5® Site-Directed Mutagenesis Kit (NEB), following manufacturer’s instructions. MEF WT and MEF *Prmt7* KO were immortalized at passage 3 by transfection of SV40LT.

### PRMT7 cellular assay

C2C12 cells were plated and next day treated with compounds. After 48 h, cells were lysed in lysis buffer (20 mM Tris-HCl pH 8, 150 mM NaCl, 1 mM EDTA, 10 mM MgCl_2_, 0.5% TritonX-100, 12.5 U.mL^-1^ benzonase (Sigma), complete EDTA-free protease inhibitor cocktail (Roche)). After 2 min incubation at RT, SDS was added to final 1% concentration. Cell lysates were analysed in western blot for unmethylated and monomethylated Hsp70/Hsc70 levels. The IC_50_ values were determined using GraphPad Prism 7 software.

### Western blot

Total cell lysates or cellular fractions (as indicated) were resolved in 4-12% Bis-Tris Protein Gels (Invitrogen) and transferred in for 1.5 h (80 V) onto PVDF membrane (Millipore) in Tris-Glycine transfer buffer containing 20% MeOH and 0.05% SDS. Blots were blocked for 1 h in blocking buffer (5% milk in PBS) and incubated with primary antibodies in blocking buffer (5% BSA in PBST: 0.1% Tween 20 PBS) overnight at 4 °C. After five washes with PBST the blots were incubated with goat-anti rabbit (IR800) and donkey anti-mouse (IR 680) antibodies in Odyssey Blocking Buffer (LiCor) for 1 h at RT and washed five times with PBST. The signal was read on an Odyssey scanner (LiCor) at 800 nm and 700 nm.

### Knock-down

Cells were transfected with 15 nM of either non-targeting siRNA or siRNA against PRMT7, PRMT4 or PRMT5 (Dharmacon) using jetPRIME® transfection reagent (Polyplus), following manufacturer instructions. After 3 days the protein levels were measured by Western blot as described above.

### Cell growth and apoptosis assay after heat shock

MEF WT and *Prmt7* KO cells were heat shocked in a water bath for 20 min at 44 °C. Cell number was determined with Vybrant® DyeCycle™ Green 24 h after heat shock and apoptosis levels were determined with IncuCyte® Caspase-3/7 Reagent within 12 h after heat shock using IncuCyte™ ZOOM live cell imaging device (Essen Bioscence) and analysed with IncuCyte™ ZOOM (2015A) software.

### Cell growth after bortezomib treatment

MEF WT and *Prmt7* KO cells were treated with bortezomib for 4 h or 24 h and the confluency monitoring was started at 4 h after bortezomib (30 nM) treatment. For experiments with SGC3027 or SGC3027N cells were pretreated with 4 μM compounds for 2 days before bortezomib treatment (30 nM). After 20 h bortezomib, SGC3027 or SGC3027N were removed and SGC3027 or SGC3027N were replaced. Cell confluency was monitored right after bortezomib addition using IncuCyte™ ZOOM live cell imaging device (Essen Bioscence) and analysed with IncuCyte™ ZOOM (2015A) software.

### Cellular fractionation

Cells were trypsinized and 1 x 10^6^ of cells were centrifuged at 400 x g for 5 min at 4 °C. Cell pellets were resuspended in 200 μL of hypotonic buffer (10 mM HEPES pH 7.5, 10 mM KCl, 1.5 mM MgCl2, 0.3 M Sucrose, 1 mM TCEP, 0.1% Triton X-100 and protease inhibitors). The cell suspensions were incubated on ice for 15 minutes followed by centrifugation at 1300 x g for 5 min at 4 °C. The supernatants were collected and cleared by centrifugation at 18,000 x g to produce the cytoplasmic fraction. The pellets were then washed in hypotonic buffer, centrifuged again and resuspended in an equal volume of lysis buffer (as described above in PRMT7 cellular assay) to produce nuclear fraction. The fractions were analysed by Western blot as described above.

### Immunofluorescence

C2C12 cells were electroporated (1650 V, 10 ms, 3 pulses) with 0.5 μg of PRMT7-FLAG plasmid using Neon transfection system (Life Technologies), following manufacturer instructions. Other cells were transfected with PRMT7-FLAG using X-tremeGeneXP (Sigma), following manufacturer instructions.The next day cells were washed with PBS, fixed with 4% PFA for 10 min, permeabilized with 0.1% Tritton X-100/PBS for 5 min, blocked with 5% BSA in PBS-T (PBS, 0.1% Tween20) for 1 h and incubated with anti-FLAG antibodies (1:1000) overnight at 4 °C. Next day cells were washed with PBS-T, incubated with anti-mouse Alexa Fluor 488 in blocking buffer (1:1000) and washed with PBS-T. Nuclei were labeled with Hoechst 33342 dye (Thermo Fisher Scientific), following manufacturer instructions. The images were taken with EVOS FL Auto 2 Imaging System (Thermo Fisher Scientific).

### Immunoprecipitation

HCT116 *PRMT7* KO (clone 94A) cells were cotransfected with GFP-tagged HSPA8/1 (WT or R469A mutant) and FLAG-tagged PRMT7 (WT or R44A mutant) at 1:10 ratio using JetPRIME transfection reagent (Polyplus), following manufacturer’s instructions. Cells were lysed in lysis buffer (20 mM Tris-HCl pH 8, 150 mM NaCl, 1 mM EDTA, 10 mM MgCl_2_, 0.1% TritonX-100 and complete EDTA-free protease inhibitor cocktail (Roche)) for 20 min and centrifuged 18,000 x g for 3 min. The supernatants were incubated with rabbit anti-GFP antibody (Invitrogen) overnight at 4 °C. Next day the antibody complexes were incubated with prewashed Dynabeads™ Protein G (Thermofisher) for 2 hours. Beads were washed in lysis buffer and proteins were eluted with 2 x SDS loading buffer and analyzed by Western blot, as described above.

### CRISPR/Cas9 gRNA vector design for HCT116 cells

For HCT116 cells 3 guide RNA were designed on PRMT7 locus. To generate gRNA expression vectors, the annealed oligonucleotide for each targeting site and annealed scaffold oligonucleotides were ligated into pENTER/U6 vector (Life Technologies). Cas9 expression vector was prepared previously ^62^. Briefly, the Cas9 cDNA was synthesized by Eurofins genomics and inserted into the pCAGGS expression vector provided by Dr. J. Miyazaki (Osaka University, Osaka, Japan) ^63^. Following guide RNA was used for generation of knockout cells: guideRNA/synthetic Oligonucleotide_sense/ synthetic Oligonucleotide_antisense (21: GGGACTCTTGTCAATGATGGCGG/caccGGGACTCTTGTCAATGATGGgtttta/ctctaaaacccatcattgacaagagtccc; 74: GGCATGGGTACTCCCACAGCGGG/caccGGCATGGGTACTCCCACAGCgtttta/ ctctaaaacgctgtgggagtacccatgcc; 94: GGGCAGCTCTCCACGTCAACGGG/ caccGGGCAGCTCTCCACGTCAACgtttta/ ctctaaaacgttgacgtggagagctgccc.

### CRISPR/Cas9-mediated genome editing

HCT116 cells were seeded onto 6-well plates at a density of 40,000 cells.well^-1^, 24 hr prior to transfection. Cells were transfected using Lipofectamine 2000 (Life Technologies) according to the manufacturer’s instruction. A total of 3 μg Cas9 expression vector, 1 μg of gRNA expression vector, and 0.4 μg of pEBMultipuro (Wako Chemicals) as a transfection marker were transfected. After 48 h of transfection, 1 μg.mL^-1^ puromycin was added for selection. The colonies were isolated by limiting dilution. PRMT7 destruction for the isolated clones was confirmed by Sanger sequencing and Western blotting.

### CRISPR for mouse C2C12 cells

C2C12 PRMT7 KO clones were generated by cloning guideRNA (caccgGTCATGTAGCATGTCGGCAT/aaacATGCCGACATGCTACATGAC) into LentiGuide-Puro vector (from Zhang lab), following Zhang laboratory protocols. Lentivirus was prodiced using standard protocols. 48 h post transfection the supernatant was collected, filtered through 0.5 μm filter and used to infect C2C12 cells in presence of polybrene to final conc of 8 μg.ml^-1^. After 24 h media was changed and after 2 days puromycin was added to final concentration of 2 μg.mL^-1^. 5 x 10^4^ puromycin selected cells were electroporated with 0.5 μg Cas9-GFP plasmid (1650 V, 10 ms, 3 pulses). Next day GFP positive cells were sorted and plated in 96-well plates (1 cell.well^-1^). The clones were analysed for PRMT7 expression and HSP70 monomethylation in western blot. The PRMT7 KO and PRMT7 catalytic mutant clones were genotyped by Sanger sequencing PCR-amplified TA-cloned genomic DNA. The premature stop codon in exon 3 or 4 resulted in PRMT7 KO. The mutation in one of the alleles delY35,A35S in addition to premature codons in exon 3 or 4 of other alleles resulted in expression of catalytically inactive PRMT7 protein (**Supplementary Table 5**)

### Cell culture for methylome analysis

*PRMT7* KO HCT116 cells (clone 94A) or parental control cells were cultivated in McCoy’s 5A medium (Gibco) containing 10% fetal bovine serum (FBS) (Hyclone), and maintained in an incubator set to 37 °C, 5% CO2. For iMethyl-SILAC-labeling of cells, DMEM without L-methionine, L-cystine and L-glutamine (Sigma) was supplemented with 0.2 mM of L-cystine and 2 mM of L-glutamine to prepare DMEM lacking only L-methionine. The L-methionine free DMEM was supplemented with 10% dialyzed fetal bovine serum (Thermofisher Scientific) and either L-methionine-^13^C_4_ (Cambridge Isotope Laboratories) or L-methionine-methyl-^13^CD_3_ (Sigma) at the final concentration of 0.1 mM. Clone 94A or control cells were labelled with medium containing L-methionine-^13^C_4_ and L-methionine-methyl-^13^CD_3_, respectively for five days.

### Enrichment of mono-methylarginine peptides

Each stable isotope labeled HCT116 cell in 10 cm dishes was washed with phosphate-buffered saline (PBS), and lysed with 0.6 mL.dish^-1^ RIPA buffer [50 mM Tris-HCl (pH 7.5), 150 mM NaCl, and 1% NP-40 containing protein phosphatase inhibitor cocktail 1 and 2 (Sigma) and protease inhibitors (Sigma)]. The cell lysate was centrifuged for 15 min at 20000 x g to remove cellular debris. The recovered proteins were quantified with a BCA Protein Assay Kit (Thermofisher Scientific). Five mg of each cell lysate was combined and precipitated with 4 volumes of cold acetone and centrifuged for 15 min at 20000 x g. The precipitates were dissolved in 8 M urea/50 mM triethylammonium bicarbonate (TEAB, Wako), reduced with 5 mM Tris(2-carboxyethyl)phosphine hydrochloride (TCEP-HCl, Thermofisher Scientific) and then digested with 100 μg of Lys-C (Wako) for 2 h at room temperature, followed by 5-fold dilution with Milli-Q water and digested with 100 μg of trypsin (Promega) overnight at room temperature. The protein digests were alkylated with iodoacetamide (IAA, Sigma) for 30 min at room temperature, acidified with 10% trifluoroacetic acid (TFA, Wako) and desalted on ODS column (C18MG 4.6 x 150 mm, Shiseido). The digested peptide mixtures were lyophilized to give white powders. Mono-methyarginine peptides were enriched by a PTMscan Mono-Methyl Arginine Motif [mme-RG] Kit (Cell Signaling) according to the manufacturer’s protocol followed by strong cation exchange (SCX) chromatographic fractionation.

### SCX fractionation

SCX chromatographic fractionation was performed on Shimazu HPLC system using a polysulfoethyl A SCX column (2.1 x 35 mm, 5 μm, 300 A, PolyLC). Mono-methyarginine peptide eluates were diluted with solvent A by 10 fold, and loaded to an equilibrated column with 80% acetonitrile (ACN) containing 0.1% HCO_2_H (Solvent A). After 10 min in 100% A, peptides were eluted with 30% ACN containing 350 mM NH_4_HCO_3_, pH 3 (Solvent B) using a gradient from 0% to 10% B in 20 min, 10% to 20% B in 10 min, 20% to 40% B in 5 min, 40% to 80% B in 5 min, followed by 80% to 100% B in 1 min and then maintained at 100% B for 30 min at a flow rate of 0.1 mL.min^-1^. Fractions were collected at 2 min interval (0.2 mL each) from 20 min to 60 min after sample injection and polled at various intervals to bring total fraction (fr) number to 8; SCX08 : fr. 1-8, SCX10 : fr. 9-10, SCX12 : fr. 11-12, SCX14 : fr. 13-14, SCX16 : fr. 15-16, SCX18 : fr. 17-18, SCX19 : fr. 19, SCX20 : fr. 20.

### LC-MS/MS analysis

The SCX column fractionated peptides were dissolved in 0.1% TFA with 2% ACN and analyzed by on-line nano LC using an Easy nLC1000 System (Thermo Fisher Scientific) coupled to Fusion Orbitrap mass spectrometer (Thermo Fisher Scientific). The peptides were loaded onto a trap column (C18 Pepmap100, 3 μm, 0.075 x 20 mm) and resolved on an analytical column (Reprosil-Pur C18AQ 3 μm 0.075 x 150 mm, Nikkyo Technos) at 300 nL.min^-1^ over 45 min. The mass spectrometer was operated in a top-20 mode with dynamic exclusion of 10 s collecting MS spectra in the Orbitrap mass analyzer at a resolution of 120000 and data-dependent collision-induced dissociation MS/MS spectra in the ion trap with normalized collision energy of 30.

### Database Search

Peak lists extraction from Xcalibur raw files were automatically performed using Proteome Discoverer 1.4 software (Thermofisher Scientific). Peak lists were searched against Uniprot using Mascot software (version 2.5, Matrix Science). The methylation level of arginine was calculated by the ratio of L/H (light/heavy) peptide peaks using Proteome Discoverer. The mass tolerance of precursor and fragment were set to 10 ppm and 0.45 Da. Up to two missed trypsin cleavages were allowed. For Mascot search, the following parameters were set: carbamidomethylation cysteine was selected as a fixed modification; oxidation of methionine, monomethylation of arginine, [2H(3)13C(1)] labeled monomethylation of arginine, [13C(4)] labeled methionine, and [2H(3)13C(1)] labeled methionine were selected as variable modifications. Peptide identification data are routinely filtered to 1% false-discovery rate using the target-decoy strategy. The LC/MS data were also searched using Peaks 8.5 software (Bioinformatics Solutions Inc.) against the human Uniprot database (downloaded on July 2018) and using the same database search parameters as above.

### Purification, crystallization and structural determination

mPRMT7 was expressed and purified according to the previously published protocol ^64^. mPRMT7 was first mixed with 5-fold molar excess of SAH and incubated for 30 min on ice. Good diffracting crystals were obtained in vapour-diffusion sitting drops, in a precipitant solution containing 0.04 M citric acid, 0.06 M Bis-Tris propane [pH 6.4], 20% (v/v) PEG 3350. The mPRMT7 crystals were soaked into a 1 μL drop containing the same precipitant solution supplemented with 1 mM SGC8158 and 20% (v/v) glycerol for 5 days at 20 °C. Crystals were then cryo-cooled in liquid nitrogen. The mPRMT7_ SGC8158 dataset was collected at the 24ID-E beamline at the Advanced Photon Source (APS). Dataset was processed with XDS ^65^ and merged with Aimless ^66^. Initial phases were obtained by using mPRMT7 (PDB ID:4C4A) as initial model in Fourier transform with refmac5 ^67^. Model building was performed in COOT ^68^ and the structure was validated with Molprobity ^69^. SGC8158 restraints were generated using Grade Web Server (http://grade.globalphasing.org). Images were prepared with PyMol Software (www.pymol.org).

### Statistical Testing

Statistical significance was assessed using GraphPad Prism 7 software via Student’s t test (unpaired, 95% confidence interval). p values < 0.05 were considered statistically significant.

### Data Availability

The mass spectrometry data have been uploaded to the ProteomeXchange Consortium via the PRIDE partner repository. The dataset accession number is PXD012119. mPRMT7_SGC8158 structure has been deposited in PDB (PDB ID: 6NPG).

