## Supplementary files for "Pharmacological inhibition of PRMT7 links arginine monomethylation to the cellular stress response"

* Equal contribution
### Corresponding authors

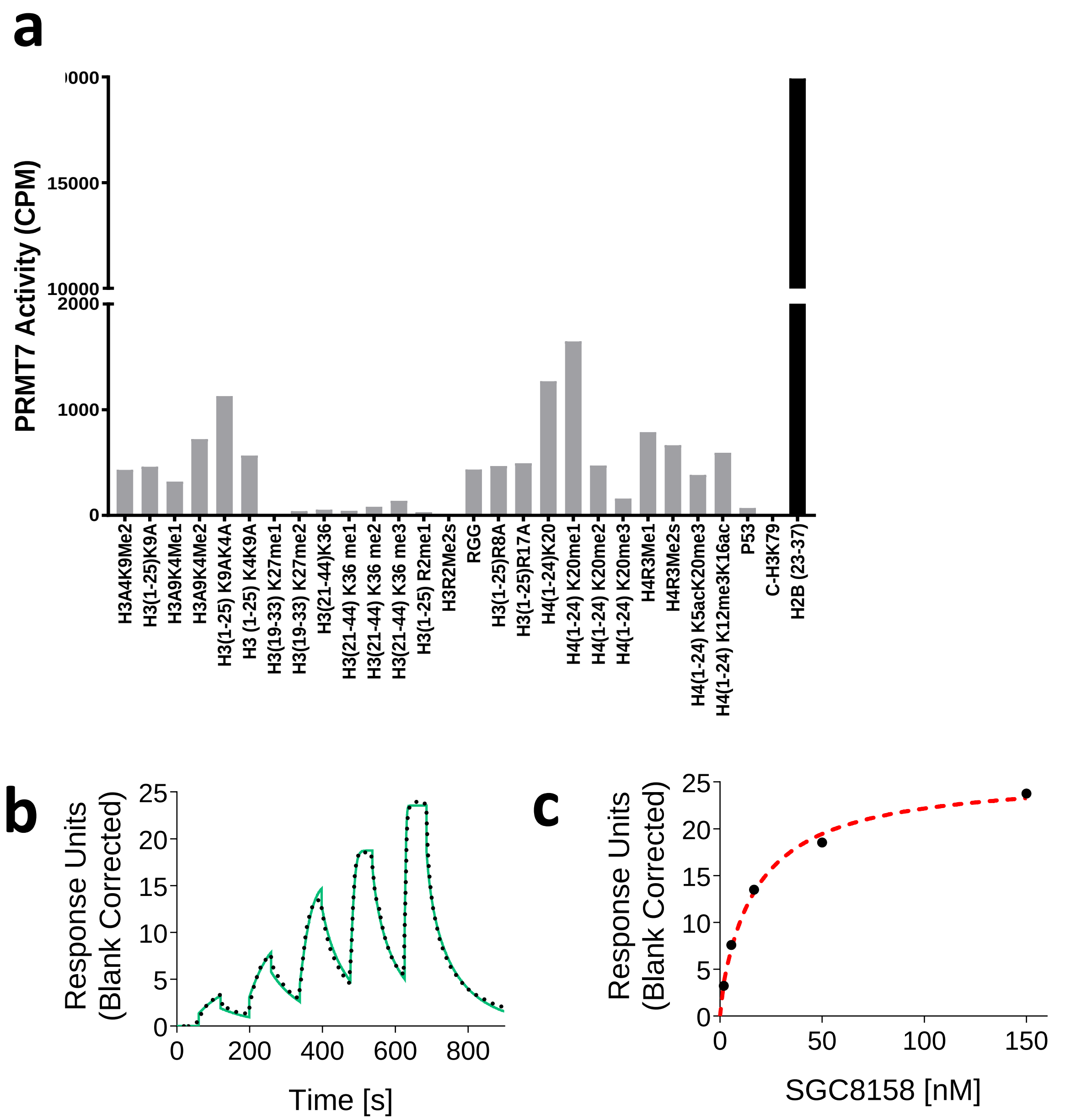

**Supplementary. Fig. 1: *In vitro* characterization of PRMT7 histone substrate specificity and SGC8158 binding. a,** PRMT7 histone substrate specificity. PRMT7 was screened against 28 biotinylated peptides with various length and modifications as potential substrates, as described in material and methods. PRMT7 was significantly active only with the H2B (23-37, KKDGKKRKRSRKESY) peptide. Data are presented as average from two independent experiments. **b,c** SPR analysis of SGC8158 binding to PRMT7. **b**, A representative SPR sensorgram (solid green) shown with the kinetic fit (black dots). $K_D$ (6.4 ± 1.2 nM), $k_{on}$ (4.4 ± 1.1 × $10^6$ $M^{-1}$ $s^{-1}$) and $k_{off}$ (2.6 ± 0.5 × $10^{-2}$ $s^{-1}$) were calculated from kinetic fitting (n=3). **c,** The steady state response (black circles) obtained from **b** with the steady state 1:1 binding model fitting (red dashed line). $K_D$ = 16.4 ± 1.0 nM (n=3).

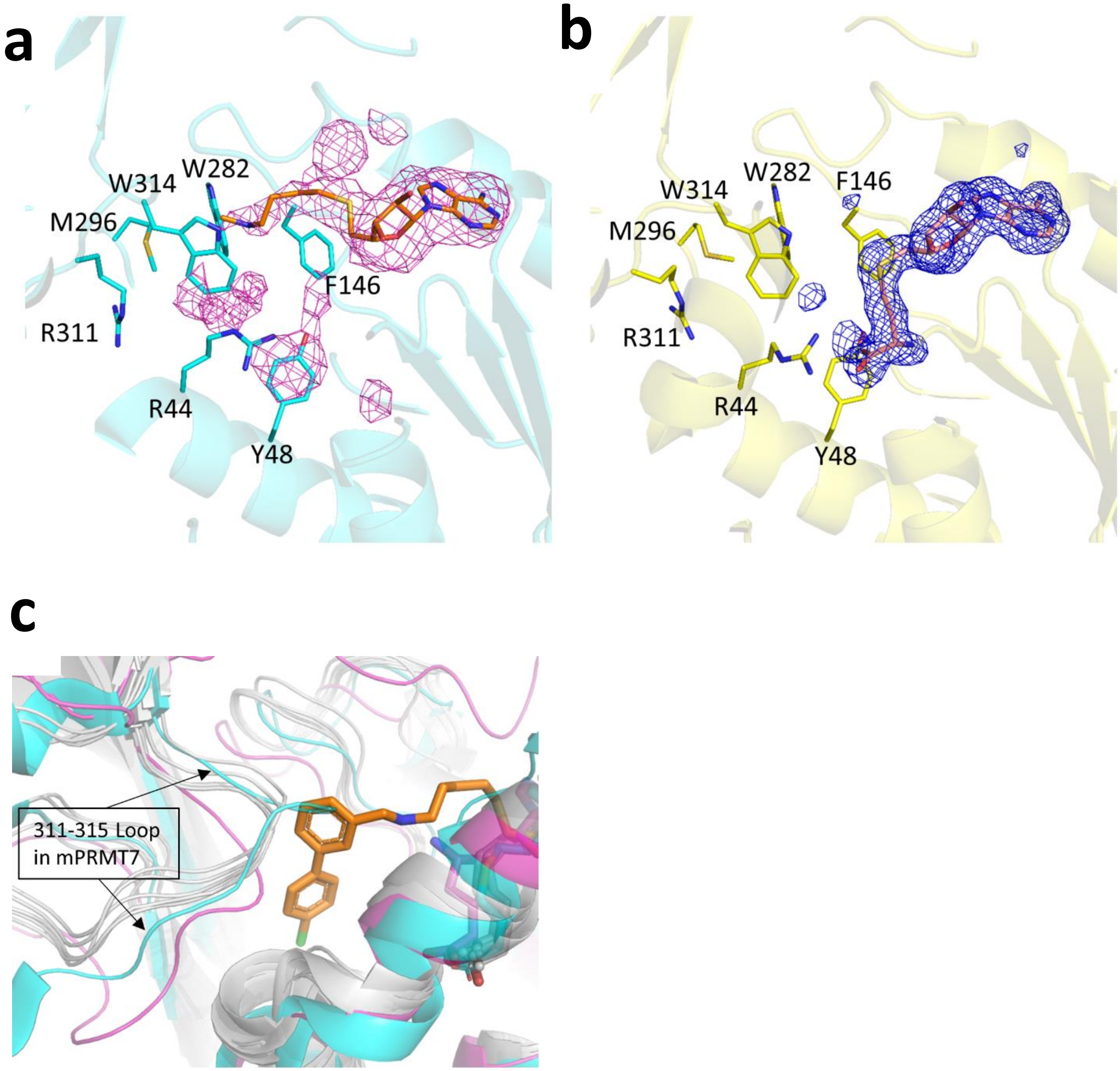

**Supplementary Figure 2: The compound binding pocket in mPRMT7_SGC8158**. **a,** The $mF_o$-$DF_c$ electron density omit-map of SGC8158 displayed as magenta mesh, contoured at 2.5σ. Only the adenosyl moiety and the linker region of SGC8158 is modeled into the density. This structure was derived from crystals in which SGC8158 was soaked into SAH-bound protein crystals to compete out the SAH. **b,** For comparison, the $mF_o$-$DF_c$ electron density omit-map of SAH in mPRMT7_SAH (PDB ID: 4C4A) is displayed as blue mesh, contoured at 3.0σ. **c,** Comparison of the 311-315 loop region in mPRMT7 (in cyan) with PRMT5 (in magenta) and PRMT1,3,4,6,8 (in grey). SGC8158 is shown as sticks model in orange.

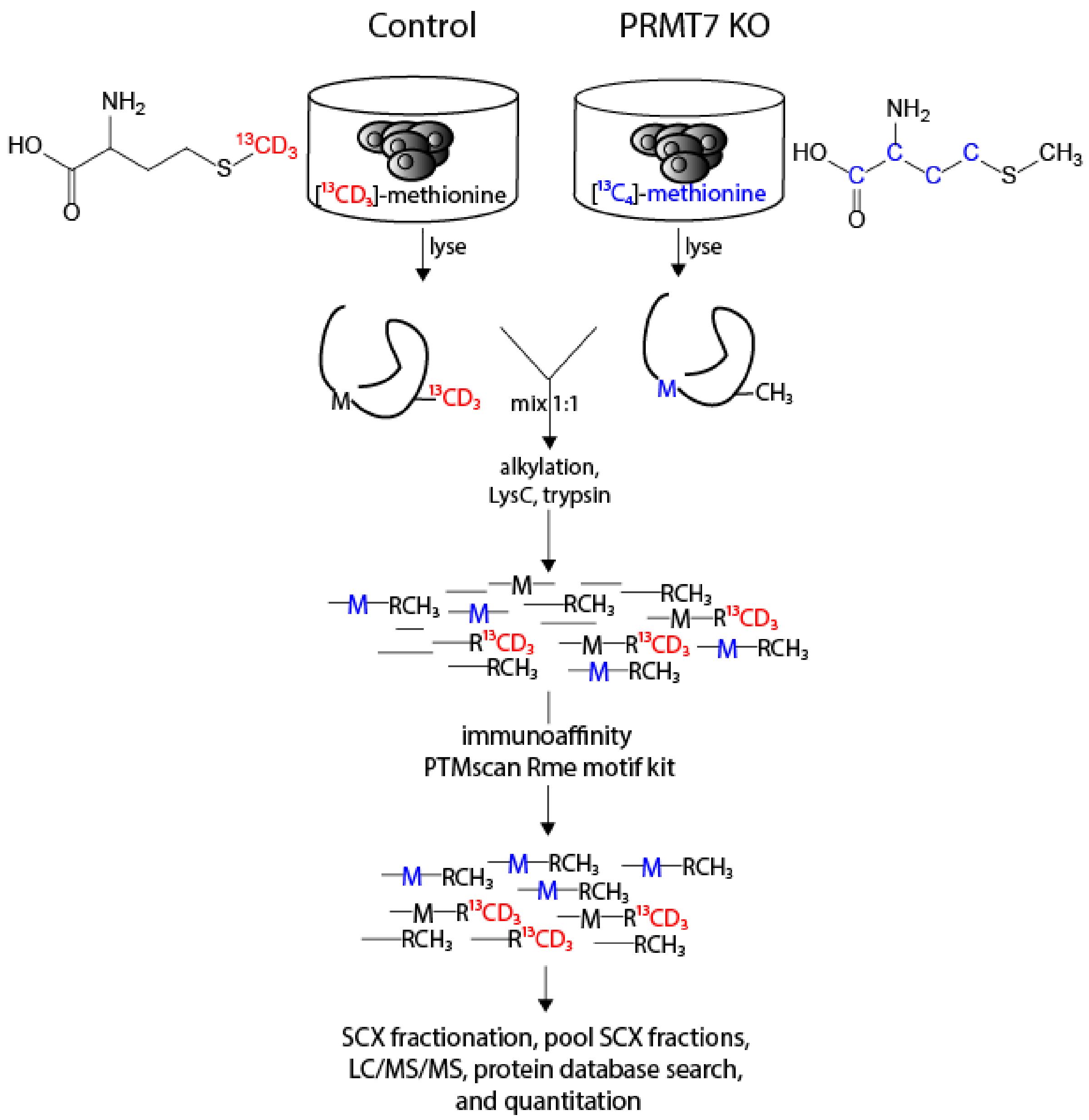

**Supplementary Fig. 3: Identification of putative PRMT7 substrates using iMethyl-SILAC.** HCT116 WT and *PRMT7* KO cells were labelled with [$^{13}CD_3$]-methionine and L-methionine-$^{13}C_4$, respectively. Protein from lysed cells were mixed at 1:1 (mass) ratio, reduced, alkylated (iodoacetamide), and digested with Lys-C followed by trypsin. Peptides containing mono-methylarginine were enriched using immunoprecipitation, fractionated by SCX chromatography and analysed by LC/MS/MS.

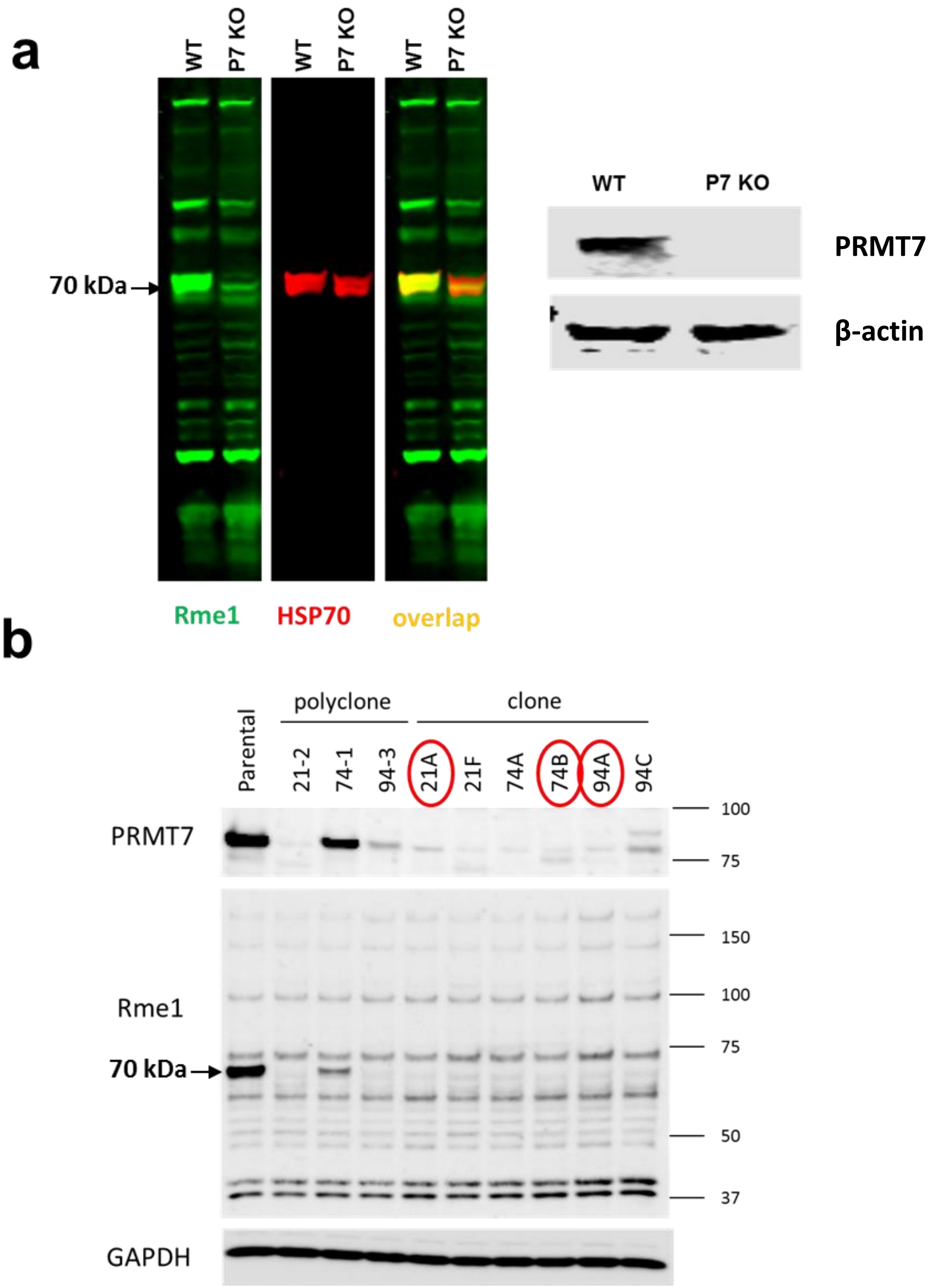

**Supplementary. Fig. 4: HSP70 family member HSPA8 as potential PRMT7 substrate. a**, Western blot analysis of *PRMT7* wild type and knockout HCT116 cell extracts (clone 94A) using monomethyl arginine antibodies (Rme1) shows decrease in 70 kDa band corresponding to HSP70 proteins in *PRMT7* KO cells. **b**, CRISPR clones analysis of *PRMT7* KO in HCT116 cells. Monomethyl arginine signal is diminished at the 70 kDa range. Polyclone cells are the ones after CRISPR/Cas9 transfection without cloning. Clones highlighted with red were used in this study.

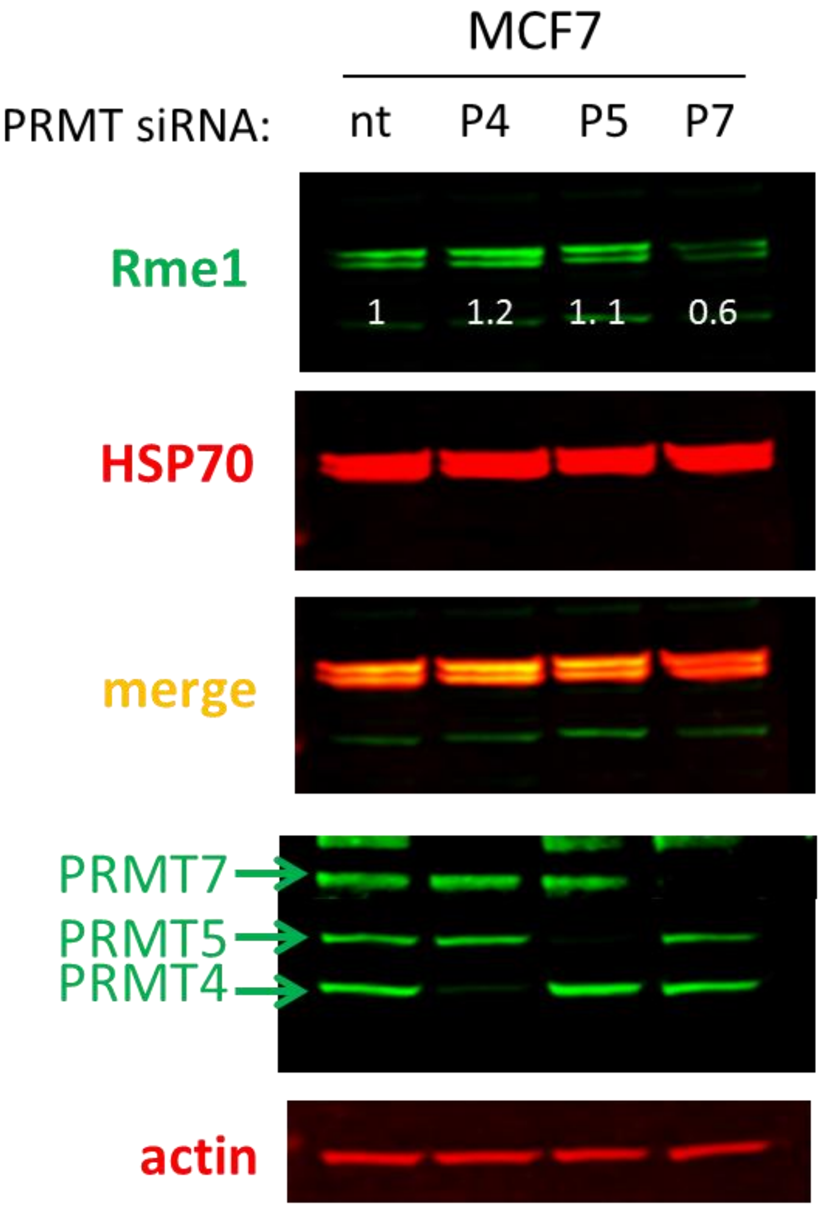

**Supplementary. Fig. 5: The knockdown of PRMT7 (P7) but not PRMT4 (P4) or PRMT5 (P5) leads to decrease in HSP70 monomethylation in MCF7 cells.** MCF7 cells were transfected with siRNA for 3 days and cytoplasmic fraction was analysed for HSP70 arginine monomethylation (Rme1) levels. **nt**- non-targeting siRNA control.

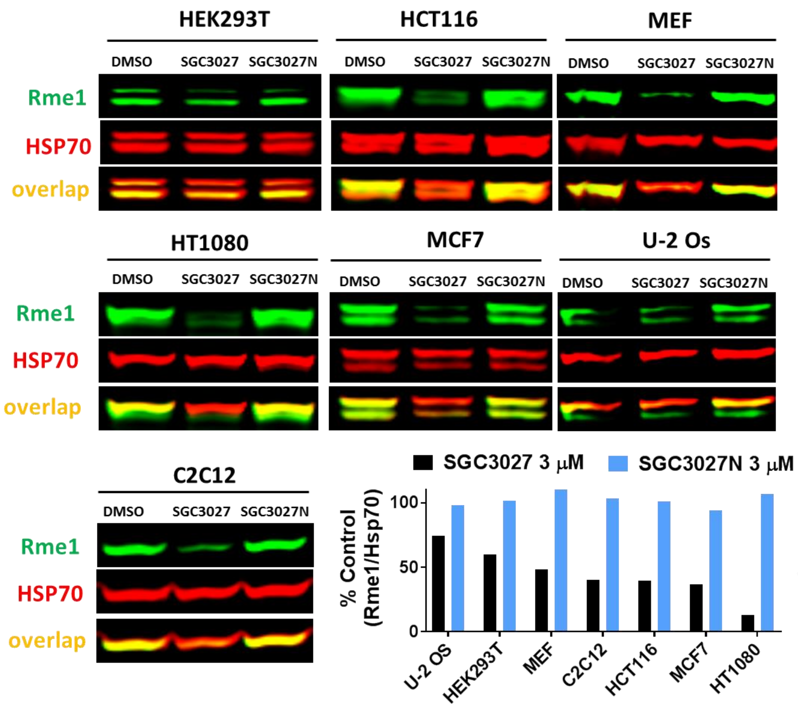

**Supplementary. Fig. 6: The effect of SGC3027 and SGC3027N on HSP70 monomethylation in different cell lines.** Cells were treated with compounds for 2 days and analysed in western blot for arginine monomethylation (Rme1) and HSP70 levels. SGC3027 cellular potency is cell line dependent. The graph represents ratios of Hsp70-Rme1 band intensities and total Hsp70 signal in cells treated with SGGC3027 and SGC3027N normalized to untreated control (n=1). Note that, in some cell lines, the pan-anti-Rme1 antibody used to detect methylation recognized other methylated proteins approximately the same size as Hsp70, thus we recommend running gels for longer time or using cytoplasmic fractions since in most cases the other methylated proteins were found in nuclear fraction.

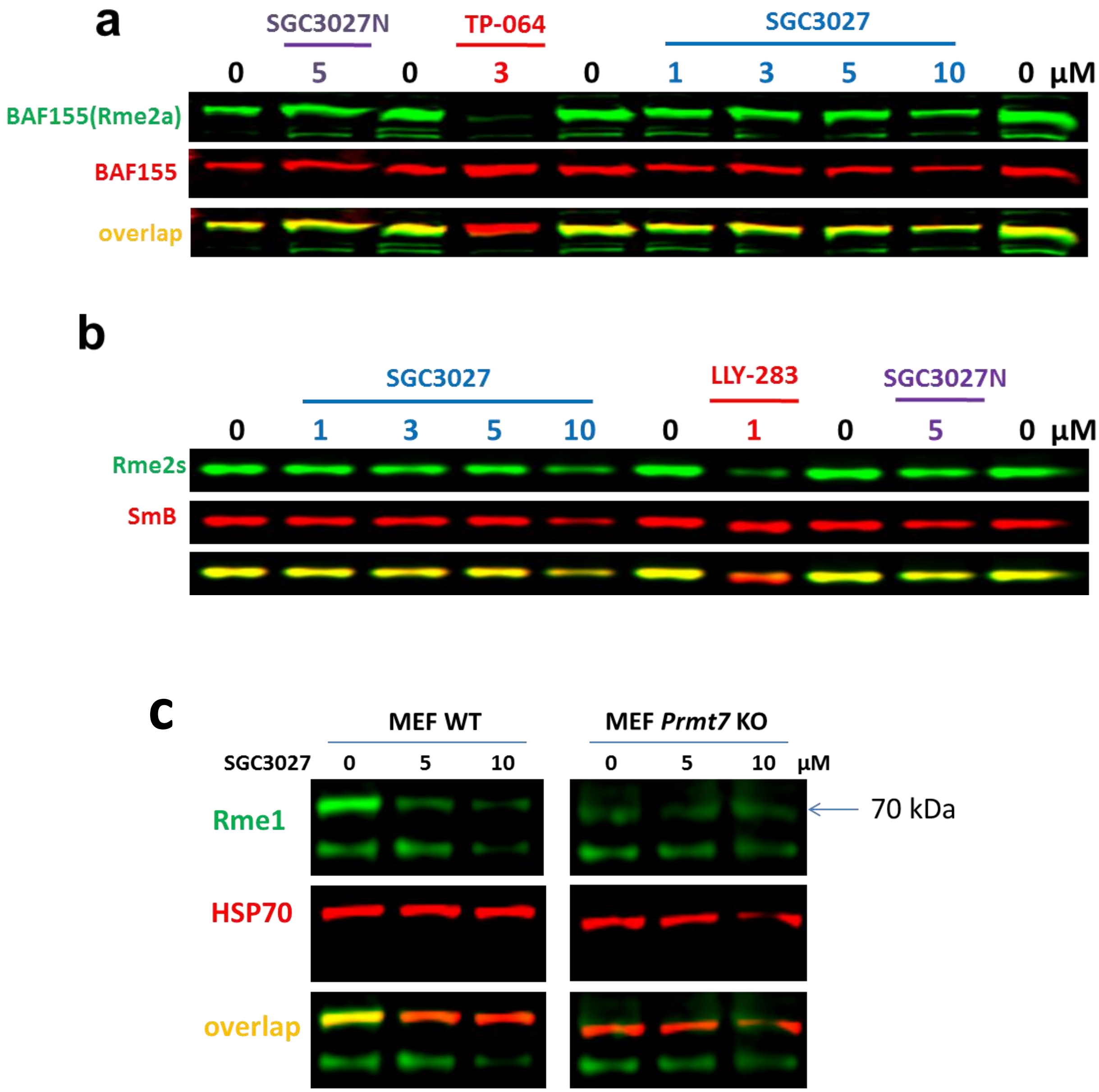

**Supplementary. Fig. 7: SGC3027 is a selective PRMT7 inhibitor in cells and does not affect the HSP70 methylation in *Prmt7* KO MEFs.** Cells were treated with compounds for 2 days. **a**, PRMT4 dependent BAF155 asymmetric arginine demethylation (Rme2a) levels were only decreased by PRMT4 selective inhibitor TP-064. **b**, PRMT5 dependent SmB symmetric arginine demethylation (Rme2s) was only decreased by PRMT5 selective inhibitor LLY-283. **c**. SGC3027 decreases Hsp70 monomethylation in MEF WT cells but not *Prmt7* KO MEFs.

**Supplementary. Table 1. Selectivity of SGC8172.**

| **Target** | **$IC_{50}$ (nM)** | **Hill Slope** |
|---|---|---|
| PRMT7 | 0.7 | -1.3 |
| PRMT1 | >5000 | NA |
| PRMT3 | 268 | -0.9 |
| PRMT4 | 4 | -1 |
| PRMT5-MEP50 | 44 | -0.7 |
| PRMT6 | ~1200 | -1.1 |
| PRMT8 | ~1500 | -1 |
| PRMT9 | 138 | -0.5 |

**Supplementary. Table 2. Selectivity of SGC8158 and SGC8158N**. Activity of 35 protein, RNA and DNA methyltransferases were assessed at 1 and 10 µM of SGC8158. The values are presented as percent activity.

| Target | Activity % | | | |
|---|---|---|---|---|
| | SGC8158 | | SGC8158N | |
| | @ 1 µM | @ 10 µM | @ 1 µM | @ 10 µM |
| **PRMT7** | 2 | 0 | 95 | 75 |
| PRMT1 | 104 | 67 | 94 | 91 |
| PRMT3 | 57 | 6 | 96 | 96 |
| PRMT4 | 7 | 0 | 101 | 100 |
| PRMT5 | 15 | 2 | 101 | 91 |
| PRMT6 | 100 | 58 | 94 | 87 |
| PRMT8 | 92 | 45 | 97 | 97 |
| PRMT9 | 19 | 4 | 106 | 95 |
| G9A | 100 | 99 | 99 | 98 |
| GLP | 99 | 95 | 95 | 90 |
| SUV39H1 | 94 | 90 | 98 | 78 |
| SUV39H2 | 103 | 102 | 103 | 78 |
| SETDB1 | 103 | 99 | 99 | 91 |
| SETD7 | 100 | 100 | 96 | 89 |
| MLL1 | 106 | 99 | 101 | 98 |
| MLL3 | 105 | 105 | 99 | 100 |
| PRDM9 | 106 | 96 | 101 | 98 |
| SETD8 | 101 | 102 | 99 | 100 |
| SUV420H1 | 96 | 87 | 99 | 102 |
| SUV420H2 | 98 | 87 | 102 | 101 |
| SETD2 | 95 | 86 | 103 | 97 |
| PRC2-EZH1 | 106 | 100 | 92 | 93 |
| PRC2-EZH2 | 105 | 94 | 93 | 88 |
| SMYD2 | 101 | 84 | 105 | 104 |
| SMYD3 | 102 | 104 | 100 | 102 |
| METTL3-14 | 97 | 84 | 87 | 73 |
| BCDIN3D | 55 | 11 | 96 | 94 |
| DNMT1 | 98 | 91 | 94 | 99 |
| DNMT3A/3L | 100 | 99 | 100 | 99 |
| DNMT3B/3L | 100 | 101 | 104 | 103 |
| NSD1 | 104 | 99 | 99 | 98 |
| NSD2 | 106 | 101 | 92 | 97 |
| NSD3 | 97 | 95 | 100 | 97 |
| DOT1L | 58 | 13 | 97 | 95 |
| ASH1L | 96 | 97 | 99 | 102 |

**Supplementary. Table 3. Evaluating the selectivity of SGC8158 for hits from Supplementary Table 2**. The effect of SGC8158 on activity of PRMTs, BCDIN3D and DOT1L were assessed by determining the $IC_{50}$ values.

| **Target** | **$IC_{50}$ (µM)** | **Hill Slope** |
|---|---|---|
| **PRMT7** | **<0.0025** | **1.1** |
| **PRMT5** | **0.12** | **0.5** |
| **PRMT4** | **0.13** | **0.8** |
| **PRMT9** | **0.13** | **0.7** |
| **BCDIN3D** | **1.1** | **0.9** |
| **PRMT3** | **1.7** | **1.6** |
| **DOT1L** | **1.7** | **1** |
| **PRMT8** | **7** | **0.7** |
| **PRMT6** | **15** | **1.2** |
| **PRMT1** | **16** | **1** |

**Supplementary. Table 4. Differentially methylated proteins identified by an iMethyl-SILAC approach with heavy (*PRMT7* WT) to light (*PRMT7* KO) ratio greater/equal than 1.3 and lower/equal than 0.77.**

| Protein Group Accessions | Gene | Ratio of Heavy/Light | | Protein Group Accessions | Gene | Ratio of Heavy/Light |
|---|---|---|---|---|---|---|
| Q709C8 | VPS13C | 2.7E+04 | | Q9Y5U2 | TSSC4 | 7.6E-01 |
| P08729 | KRT7 | 1.5E+04 | | P52597 | HNRNPF | 7.5E-01 |
| Q92945 | KHSRP | 1.3E+04 | | Q13151 | HNRNPA0 | 7.5E-01 |
| P52272 | HNRNPM | 7.5E+03 | | Q14527 | HLTF | 7.5E-01 |
| O95793 | STAU1 | 6.1E+03 | | Q9P2N5 | RBM27 | 7.4E-01 |
| P11142 | HSPA8 | 6.1E+03 | | Q96L91 | EP400 | 7.4E-01 |
| Q7Z6Z7 | HUWE1 | 5.2E+03 | | O43896 | KIF1C | 7.4E-01 |
| P37802 | TAGLN2 | 5.1E+03 | | Q92734 | TFG | 7.3E-01 |
| P67809 | YBX1 | 4.1E+03 | | Q9HAH7 | FBRS | 7.2E-01 |
| Q9Y520 | PRRC2C | 4.0E+03 | | P46783 | RPS10 | 7.1E-01 |
| Q07666 | KHDRBS1 | 4.0E+03 | | Q8NFW8 | CMAS | 7.1E-01 |
| P05783 | KRT18 | 2.0E+03 | | Q8WXE0 | CASKIN2 | 7.1E-01 |
| O43823 | AKAP8 | 2.0E+03 | | P49848 | TAF6 | 7.1E-01 |
| P08107 | HSPA6 | 4.3E+01 | | O00268 | TAF4 | 7.1E-01 |
| Q9UN86 | G3BP2 | 2.5E+01 | | Q9BX40 | LSM14B | 7.0E-01 |
| Q9NRF2 | SH2B1 | 1.0E+01 | | P61960 | UFM1 | 7.0E-01 |
| P05787 | KRT8 | 5.5E+00 | | Q9HBM6 | TAF9B | 7.0E-01 |
| Q9Y3X0 | CCDC9 | 3.4E+00 | | Q86US8 | SMG6 | 7.0E-01 |
| O15446 | CD3EAP | 2.4E+00 | | C9JLR9 | C11orf95 | 6.9E-01 |
| Q86YP4 | GATAD2A | 2.3E+00 | | O60506 | SYNCRIP | 6.8E-01 |
| P49247 | RPIA | 1.9E+00 | | Q8NBF6 | AVL9 | 6.8E-01 |
| Q71F56 | MED13L | 1.9E+00 | | Q6EEV4 | POLR2M | 6.8E-01 |
| P31942 | HNRNPH3 | 1.8E+00 | | Q86V81 | ALYREF | 6.7E-01 |
| Q9P2D1 | CHD7 | 1.8E+00 | | Q12888 | TP53BP1 | 6.6E-01 |
| Q9BST9 | RTKN | 1.7E+00 | | Q01201 | RELB | 6.5E-01 |
| P51991 | HNRNPA3 | 1.6E+00 | | Q13243 | SRSF5 | 6.5E-01 |
| Q13283 | G3BP1 | 1.6E+00 | | Q5TGY3 | AHDC1 | 6.4E-01 |
| Q96QC0 | PPP1R10 | 1.6E+00 | | P52824 | DGKQ | 6.4E-01 |
| P78344 | EIF4G2 | 1.6E+00 | | O15320 | CTAGE5 | 6.2E-01 |
| Q9Y446 | PKP3 | 1.5E+00 | | P53621 | COPA | 6.1E-01 |
| P11940 | PABPC1 | 1.5E+00 | | Q9Y2S6 | TMA7 | 6.1E-01 |
| Q9BTC0 | DIDO1 | 1.5E+00 | | Q8NFD5 | ARID1B | 5.8E-01 |
| O00267 | SUPT5H | 1.5E+00 | | Q9P1Y5 | CAMSAP3 | 5.2E-01 |
| Q14103 | HNRNPD | 1.4E+00 | | Q86W92 | PPFIBP1 | 4.5E-01 |
| P08727 | KRT19 | 1.4E+00 | | P35900 | KRT20 | 3.8E-01 |
| P17844 | DDX5 | 1.3E+00 | | Q5SW79 | CEP170 | 1.2E-01 |
| Q14444 | CAPRIN1 | 1.3E+00 | | P04792 | HSPB1 | 1.0E-01 |
| Q9H307 | PNN | 1.3E+00 | | P68366 | TUBA4A | 4.0E-02 |
| | | | | Q13232 | NME3 | 4.1E-04 |
| | | | | O95613 | PCNT | 2.2E-05 |

**Supplementary. Table 5. Characterization of PRMT7 catalytic mutant clone**. Clone 32 was genotyped by Sanger sequencing PCR-amplified TA-cloned genomic DNA. We have found three mutations in clone 32 genome which is consistent with near tetraploid karyotype of C2C12[1].

| | **Exon3 mutation** | **Exon3 and 4 aminoacid sequence** | **Type of mutation** |
|---|---|---|---|
| PRMT7 | | SSYADMLHDKDRNIKYYQGIRAAVSRVKDRGQKALVLDIGTGTG LLSMMAVTAGADFCYAIE | |
| 32-1 | DelCCGAC | SSYDAT-QRQKYKILPGYPGSCEQGERQRTEGLGS-HWHWHRPLVNDGSYCRG-LLLCYR | premature stop codon in exon 3 |
| 32-2 | DelATG | SSSDMLHDKDRNIKYYQGIRAAVSRVKDRGQKALVLDIGTGTGLL SMMAVTAGADFCYAIE | Y35del and A36S mutation |
| 32-3 | DelCCGAC | SSYDAT-QRQKYKILPGYPGSCEQGERQRTEGLGS-HWHWHRPLVNDGSYCRG-LLLCYR | premature stop codon in exon 3 |
| 32-4 | DelCCGAC | SSYDAT-QRQKYKILPGYPGSCEQGERQRTEGLGS-HWHWHRPLVNDGSYCRG-LLLCYR | premature stop codon in exon 3 |
| 32-5 | DelTATG | SSPTCYMTKTEI-NTTRVSGQL-AG-KTEDRRPWFLTLALAQASCQ-WQLLQGLTSAMLS | premature stop codon in exon 4 |
| 32-6 | DelTATG | SSPTCYMTKTEI-NTTRVSGQL-AG-KTEDRRPWFLTLALAQASCQ-WQLLQGLTSAMLS | premature stop codon in exon 4 |
| 32-7 | DelATG | SSSDMLHDKDRNIKYYQGIRAAVSRVKDRGQKALVLDIGTGTGLL SMMAVTAGADFCYAIE | Y35del and A36S mutation |
| 32-8 | Del ATG | SSSDMLHDKDRNIKYYQGIRAAVSRVKDRGQKALVLDIGTGTGLL SMMAVTAGADFCYAIE | Y35del and A36S mutation |
| 32-9 | DelTATG | SSPTCYMTKTEI-NTTRVSGQL-AG-KTEDRRPWFLTLALAQASCQ-WQLLQGLTSAMLS | premature stop codon in exon 4 |

**Supplementary Table 6. Data collection and refinement statistics**

| | mPRMT7_SGC8158 (PDB ID: 6NPG) |
|---|---|
| **Data collection** | |
| Space group | $P4_32_12$ |
| Cell dimensions | |
| *a*, *b*, *c* (Å) | 98.2,98.2,170.3 |
| α, β, γ (°) | 90.0,90.0,90.0 |
| Resolution (Å) | 50.0 - 2.94 |
| $R_{sym}$ or $R_{merge}$ | 0.103 |
| $I / \sigma I$ | 2.03 |
| Completeness (%) | 99.7 |
| Redundancy | 6.8 |
| **Refinement** | |
| Resolution (Å) | 50.0 - 2.94 |
| No. reflections | 17351 |
| $R_{work} / R_{free}$ | 18.0/25.6 |
| No. atoms | |
| Protein | 4784 |
| Ligand/ion | 25 |
| Water | 4 |
| *B*-factors | |
| Protein | 72.8 |
| Ligand/ion | 63.7 |
| Water | 46.1 |
| R.m.s. deviations | |
| Bond lengths (Å) | 0.007 |
| Bond angles (°) | 1.232 |

*Values in parentheses are for highest-resolution shell.

**PRMT7 chemical probe synthesis: Experimental Procedures and Characterization data**

**General Considerations.** Reactions were carried out under nitrogen or argon atmosphere with dry solvents using anhydrous conditions unless otherwise stated. All solvents were purchased from commercial sources and used as received. All fine chemicals were obtained from Sigma-Aldrich, or Combi-Blocks and used without further purification unless otherwise stated. Yields refer to chromatographically and spectroscopically ($^1$H-NMR) homogeneous materials, unless otherwise stated. NMR spectra were recorded on a Bruker AV-500 spectrometer and calibrated using residual undeuterated solvent as an internal reference ($CHCl_3$ @ δ 7.26 ppm $^1$H NMR, MeOD-d4 @ δ 3.31 ppm $^1$H NMR).

**Scheme 1.** Synthesis of the PRMT7 chemical probe

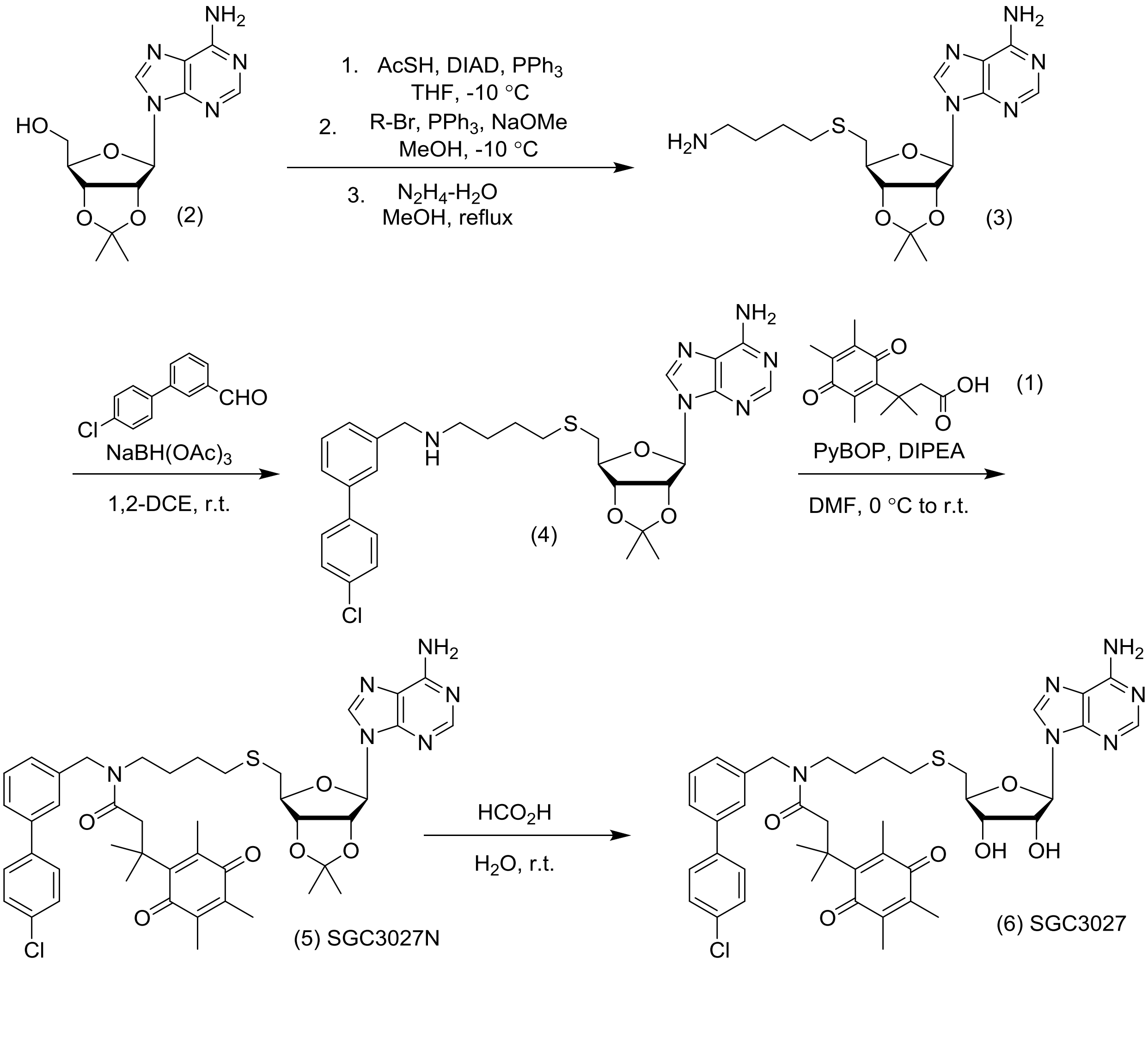

**6-hydroxy-4,4,5,7,8-pentamethylchroman-2-one**

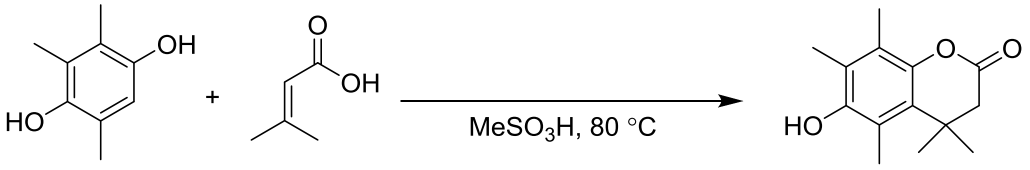

A solution of 3,3-dimethylacrylic acid (1.97 g, 19.7 mmol) and trimethylhydroquinone (3.00 g, 19.7 mmol) in methanesulfonic acid (5 mL) was heated at 70 °C in an oil bath with stirring for 2 h. The mixture was poured into water (100 mL) and extracted with EtOAc (3 x 25 mL). The combined extracts were washed with water (50 mL), saturated $NaHCO_3$ solution (2 x 50 mL), and saturated NaCl solution (50 mL) then dried over $Na_2SO_4$ prior to removal of solvent under reduced pressure to give 6-hydroxy-4,4,5,7,8-pentamethylchroman-2-one (4.47 g, 97 % yield) as a white crystalline solid.

$^1$H NMR (500 MHz, $CDCl_3$) δ 4.61 (d, *J* = 2.4 Hz, 1H), 2.55 (s, 2H), 2.36 (s, 3H), 2.22 (s, 3H), 2.18 (s, 3H), 1.45 (s, 6H).

LCMS HSS RT = 1.81 min, $[M+1]^+$ = 235.2, Purity (UV254) = 99%. Conforms to desired product.

**3-methyl-3-(2,4,5-trimethyl-3,6-dioxocyclohexa-1,4-dien-1-yl)butanoic acid (1)**

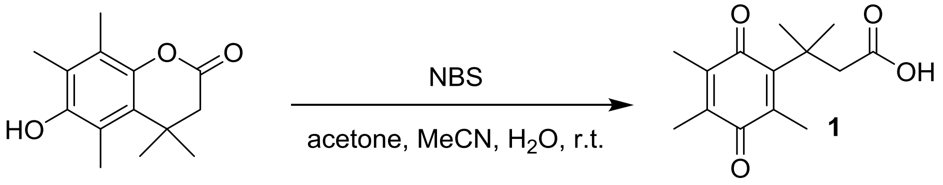

To a solution of 6-hydroxy-4,4,5,7,8-pentamethylchroman-2-one (0.501 g, 2.13 mmol) dissolved in a mixture of acetonitrile (18 mL), acetone (4 mL), and water (18 mL) was added N-bromosuccinimide (0.418 g, 2.347 mmol) in three portions and the mixture was stirred at room temperature for 30 min. After removal of the organic solvents under a stream of nitrogen overnight, the remaining yellow crystalline solid suspended in water was collected by vacuum filtration and washed with water (2 x 20 mL) to give **1** (0.495 g, 93 % yield) as a yellow crystalline solid.

$^1$H NMR (500 MHz, $CDCl_3$) δ 3.02 (s, 2H), 2.14 (s, 3H), 1.95 (s, 3H), 1.93 (s, 3H), 1.43 (s, 6H).
LCMS HSS RT = 1.75 min, $[M+1]^+$ = 251.5, Purity (UV254) = 99%. Conforms to desired product

**9-((3a*R*,4*R*,6*S*,6a*S*)-6-(((4-aminobutyl)thio)methyl)-2,2-dimethyltetrahydrofuro[3,4-d][1,3]dioxol-4-yl)-9H-purin-6-amine (3)**

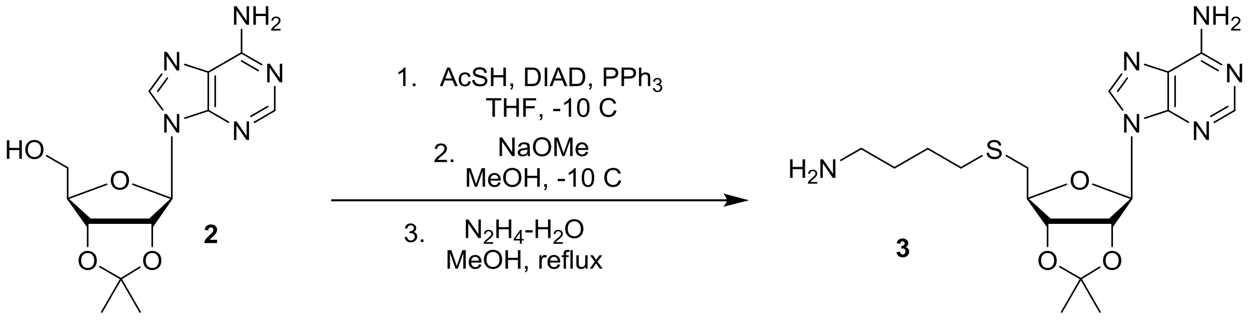

To triphenylphosphine (9.39 g, 35.8 mmol) in THF at –10 °C was added diisopropyl azodicarboxylate (7.05 mL, 35.8 mmol) over 15 minutes and the resulting slurry stirred vigorously. After 20 minutes at this temperature, 2',3'-O-isopropylideneadenosine (5.00 g, 16.3 mmol) was added as a solid in one portion and the reaction mixture stirred for an hour prior to addition of thioacetic acid (2.56 mL, 35.8 mmol). The reaction mixture was allowed to stir for another hour prior to concentration under reduced pressure. The crude product was then column chromatographed (silica gel, $CH_2Cl_2$/MeOH, 100:0 to 95:5 v/v) to afford intermediate thioester (4.88 g, 82% yield) as a colorless oil. The product was adulterated with triphenylphosphine oxide but this does not affect subsequent reactions.

LCMS HSS RT = 1.39 min, $[M+1]^+$ = 365.9, Purity (UV254) = >95%. Conforms to desired product.

To the thioester (4.02 g, 11.0 mmol) produced above in MeOH (40 mL) were added N-(4-bromobutyl)phthalimide (4.04 g, 14.3 mmol), triphenylphosphine (0.290 g, 1.10 mmol), and DMF (5 mL) prior to cooling the solution to – 10 °C. After stirring at this temperature for 15 minutes, NaOMe (30% w/v in MeOH, 4.08 mL, 22.0 mmol) was then added. The reaction was then allowed to warm to room temperature over 1 h and stirred at this temperature for 5 h at which point alkylation was complete and no starting material remained. The reaction mixture was then further diluted with MeOH (50 mL), hydrazine monohydrate (5.34 mL, 110 mmol) added and the solution heated to reflux for 30 minutes at which point LCMS showed complete deprotection of the phthalimide protecting group. Upon cooling to room temperature, Celite was added and all volatiles removed under reduced pressure. The dry-loaded product was then purified by column chromatography (RP-C18, $H_2O$ (0.5% $NH_4OH$)/MeCN, 98:2 to 10:90 v/v) to afford **(3)** (2.91 g, 67% yield) as a white powder.

$^1$H NMR (500 MHz, MeOD-$d_4$) δ 8.30 (s, 1H), 8.25 (s, 1H), 6.20 (d, J = 2.0 Hz, 1H), 5.57 (dd, J = 6.3, 2.1 Hz, 1H), 5.08 (dd, J = 6.2, 2.8 Hz, 1H), 4.36 (td, J = 6.8, 2.8 Hz, 1H), 2.80 (d, J = 6.9 Hz, 2H), 2.57 (t, J = 6.7 Hz, 2H), 2.49 (t, J = 6.8 Hz, 2H), 1.60 (s, 3H), 1.55 – 1.42 (m, 4H), 1.41 (s, 3H).

LCMS HSS RT = 1.26 min, $[M+1]^+$ = 395.5, Purity (UV254) = >95%. Conforms to desired product

**9-((3a*R*,4*R*,6*S*,6a*S*)-6-(((4-(((4'-chloro-[1,1'-biphenyl]-3-yl)methyl)amino)butyl)thio)methyl)-2,2-dimethyltetrahydrofuro[3,4-d][1,3]dioxol-4-yl)-9H-purin-6-amine (4)**

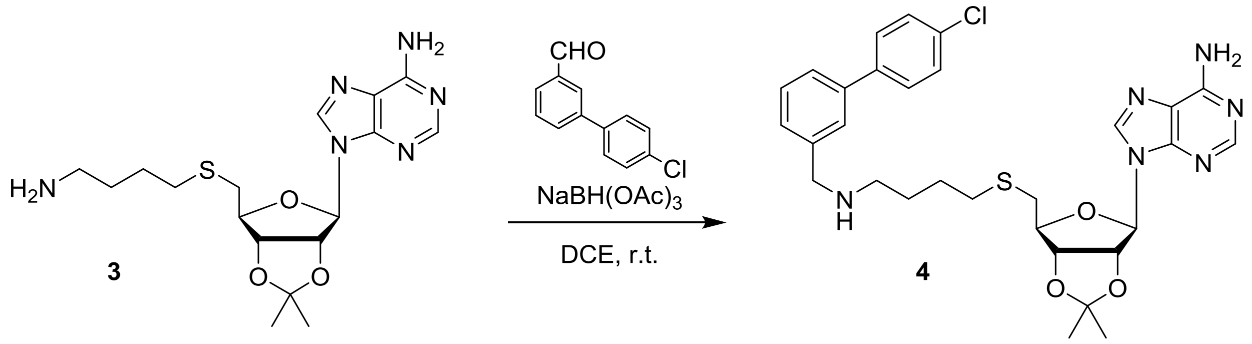

To a solution of **(3)** (115 mg, 0.292 mmol) and 4'-chloro-[1,1'-biphenyl]-3-carbaldehyde (63.2 mg, 0.292 mmol) in 1,2-dichloroethane (10 mL) was added sodium triacetoxyborohydride (93.0 mg, 0.437 mmol) in one portion at room temperature. The mixture was stirred for 2 h prior to addition of a saturated solution of aqueous $NaHCO_3$ (20 mL). The mixture was extracted with EtOAc (3 x 10 mL) and the combined organic layers dried over $Na_2SO_4$ prior to concentration under reduced pressure to afford a crude oil which was column chromatographed (silica gel, $CH_2Cl_2$/MeOH/$NH_4OH$, 100:0:0 to 96.7:3:0.3 v/v) to give **(4)** (0.142 g, 82 % yield) as a colourless semi-solid.

$^1$H NMR (500 MHz, $CDCl_3$) δ 8.35 (s, 1H), 7.94 (s, 1H), 7.56 (s, 1H), 7.53 (d, *J* = 8.5 Hz, 2H), 7.45 (d, *J* = 7.6 Hz, 1H), 7.42 – 7.37 (m, 3H), 7.33 (d, *J* = 7.3 Hz, 1H), 6.07 (d, *J* = 2.1 Hz, 1H), 5.62 (s, 2H), 5.52 (dd, *J* = 6.4, 2.1 Hz, 1H), 5.04 (dd, *J* = 6.4, 3.0 Hz, 1H), 4.38 (td, *J* = 6.8, 3.0 Hz, 1H), 3.87 (s, 2H), 2.80 (dd, *J* = 13.6, 7.1 Hz, 1H), 2.74 (dd, *J* = 13.6, 6.6 Hz, 1H), 2.65 (t, *J* = 7.0 Hz, 2H), 2.47 (t, *J* = 6.6 Hz, 2H), 1.64 – 1.51 (m, 7H), 1.39 (s, 3H); LCMS HSS RT = 1.58 min, $[M+1]^+$ = 595.7, Purity (UV254) = 99%. Conforms to desired product.

**N-(4-((((3aS,4S,6R,6aR)-6-(6-amino-9H-purin-9-yl)-2,2-dimethyltetrahydrofuro[3,4-d][1,3]dioxol-4-yl)methyl)thio)butyl)-N-((4'-chloro-[1,1'-biphenyl]-3-yl)methyl)-3-methyl-3-(2,4,5-trimethyl-3,6-dioxocyclohexa-1,4-dien-1-yl)butanamide (5) (SGC3027N)**

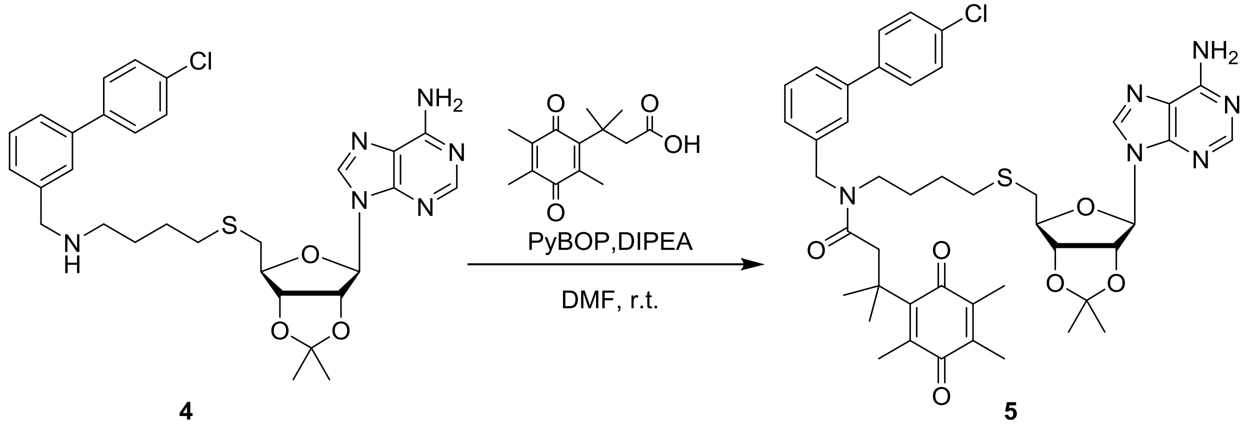

4 5

To a solution of **(4)** (62.0 mg, 0.104 mmol) in N,N-Dimethylformamide (DMF) (1 mL) were added 3-methyl-3-(2,4,5-trimethyl-3,6-dioxocyclohexa-1,4-dien-1-yl)butanoic acid (26.1 mg, 0.104 mmol) and triethylamine (36 µL, 0.260 mmol). The mixture was cooled to 0 °C prior to addition of PyBOP (81 mg, 0.156 mmol) and the mixture stirred for 3 h at this temperature. The reaction was quenched with a saturated solution of brine (20 mL) and the aqueous phase extracted with EtOAc (3 x 10 mL). The combined organic layers were dried over $Na_2SO_4$, concentrated under reduced pressure and the yellow residue column chromatographed (silica gel, $CH_2Cl_2$/MeOH/$NH_4Cl$, 100:0:0 to 95.6:4:0.4 v/v) to afford **SGC3027N** (**5)** (66 mg, 77 % yield) as a bright yellow oil (mixture of isomers).

$^1$H NMR (500 MHz, $CDCl_3$) δ 8.33 (d, *J* = 4.2 Hz, 1H), 7.91 (d, *J* = 1.8 Hz, 1H), 7.59 (d, *J* = 8.5 Hz, 1H), 7.52 – 7.44 (m, 2H), 7.44 – 7.38 (m, 3H), 7.38 – 7.31 (m, 1H), 7.22 – 7.09 (m, 1H), 6.07 (dd, *J* = 7.8, 2.0 Hz, 1H), 5.99 (d, *J* = 12.1 Hz, 2H), 5.52 (ddd, *J* = 15.4, 6.4, 2.0 Hz, 1H), 5.05 (ddd, *J* = 19.8, 6.4, 3.1 Hz, 1H), 4.55 (d, *J* = 11.9 Hz, 2H), 4.42 – 4.31 (m, 1H), 3.25 – 3.16 (m, 2H), 3.09 (d, *J* = 17.2 Hz, 2H), 2.87 – 2.67 (m, 2H), 2.54 – 2.44 (m, 2H), 2.17 – 2.10 (m, 3H), 1.95 – 1.75 (m, 6H), 1.67 – 1.60 (m, 4H), 1.56 – 1.49 (m, 2H), 1.48 – 1.42 (m, 4H), 1.41 – 1.37 (m, 6H). $^{13}$C NMR (126 MHz, $CDCl_3$) δ 191.43, 187.84, 172.37, 155.67, 154.87, 153.31, 149.43, 143.32, 140.93, 140.21, 139.49, 138.48, 137.57, 136.11, 133.89, 129.62, 129.13, 128.59, 126.34, 125.74, 124.88, 120.50, 114.58, 91.06, 86.94, 84.16, 83.96, 51.07, 46.42, 37.65, 34.49, 32.44, 28.68, 27.24, 26.74, 26.59, 25.48, 14.30, 12.60, 12.25. Complicated by rotamers (major rotamer reported); LCMS HSS RT = 2.59 min, $[M+1]^+$ = 828.0, Purity (UV254) = 99%. Conforms to desired product.

**N-(4-((((2S,3S,4R,5R)-5-(6-amino-9H-purin-9-yl)-3,4-dihydroxytetrahydrofuran-2-yl)methyl)thio)butyl)-N-((4'-chloro-[1,1'-biphenyl]-3-yl)methyl)-3-methyl-3-(2,4,5-trimethyl-3,6-dioxocyclohexa-1,4-dien-1-yl)butanamide (6) (SGC3027)**

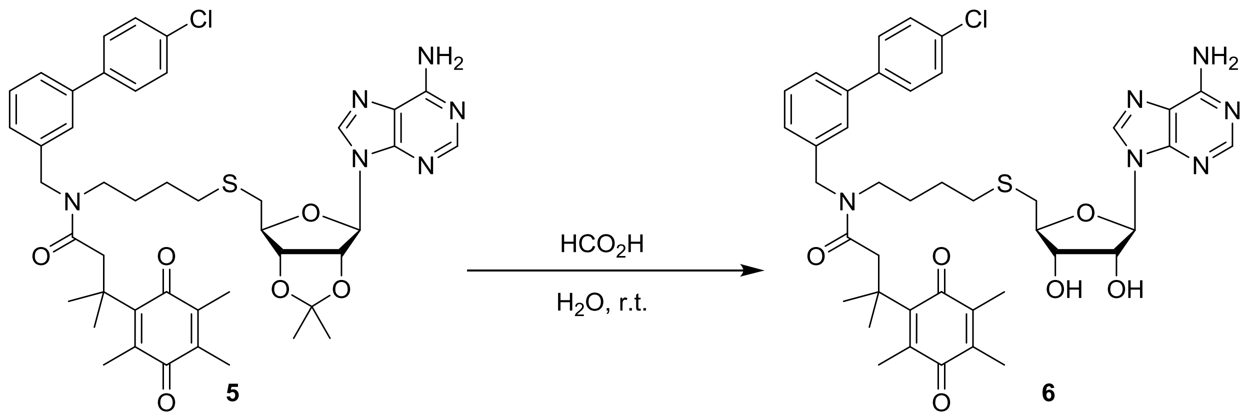

A solution of **(5)** (25.7 mg, 0.031 mmol) in $HCO_2H/H_2O$ (2 mL, 4:1 v/v) was stirred overnight at room temperature. All volatiles were then evaporated and the crude yellow residue column chromatographed (silica gel, $CH_2Cl_2$/MeOH/$NH_4OH$, 100:0:0 to 89:10:1 v/v) to afford **SGC3027 (6)** (21 mg, 86 % yield) as a bright yellow oil which solidified on standing (mixture of rotamers).

$^1$H NMR (500 MHz, $CDCl_3$) δ 8.18 (s, 1H), 8.00 (d, *J* = 1.9 Hz, 1H), 7.57 (d, *J* = 8.5 Hz, 1H), 7.51 – 7.42 (m, 2H), 7.41 – 7.29 (m, 4H), 7.19 – 7.07 (m, 1H), 6.22 – 6.08 (m, 2H), 5.97 – 5.90 (m, 1H), 4.72 – 4.63 (m, 1H), 4.61 – 4.50 (m, 2H), 4.40 – 4.30 (m, 2H), 3.30 – 3.18 (m, 2H), 3.13 – 3.04 (m, 2H), 2.90 – 2.74 (m, 2H), 2.61 – 2.49 (m, 2H), 2.11 (s, 3H), 1.93 – 1.73 (m, 6H), 1.71 – 1.63 (m, 1H), 1.60 – 1.46 (m, 3H), 1.44 – 1.34 (m, 6H); $^{13}$C NMR (126 MHz, $CDCl_3$) δ 191.38, 187.89, 172.55, 155.65, 154.62, 152.75, 149.32, 143.17, 140.95, 139.18, 138.38, 137.42, 136.47, 133.90, 129.65, 129.14, 128.58, 126.92, 126.47, 125.73, 124.89, 120.09, 90.06, 85.04, 75.00, 73.25, 53.55, 51.12, 47.78, 45.56, 37.72, 32.76, 28.78, 26.87, 14.38, 12.87, 12.28; LCMS HSS RT = 2.22 min, $[M+1]^+$ = 787.7, Purity (UV254) = 99%. Conforms to desired product.
